# Mutagenic and carcinogenic potency determinations for NDMA support the cumulative dose assumption underpinning the less-than-lifetime Threshold of Toxicological Concern

**DOI:** 10.64898/2026.01.27.702058

**Authors:** John W. Wills, Angela White, Danielle S. G. Harte, Ruby Buckley, James S. Harvey, Anthony M. Lynch

## Abstract

The management of *N-*nitrosamine impurities challenges pharmaceutical development and regulation worldwide. Because most medicinal exposures are shorter than lifetime and absolute impurity exclusion is impossible, reliable approaches to define duration-specific intake limits are essential. On the premise that carcinogenic risk is proportional to cumulative dose, the ‘Less-Than-Lifetime’ (LTL) Threshold of Toxicological Concern (TTC) framework defines progressively lower intake limits for mutagenic impurities over longer exposures. However, *N-*nitrosamines are currently treated as a ‘cohort of concern’, necessitating compound-specific evaluation placing reliance on *in vivo* mutagenicity assays for impurity qualifications. To better understand durational potency relationships and the application domain of the LTL-TTC, we apply benchmark dose (BMD) modelling to cumulative-dose-scaled transgenic rodent (TGR), error-corrected sequencing and rodent carcinogenicity datasets for *N-*nitrosodimethylamine (NDMA) obtained from the published literature. For TGR, cumulative-dose scaling better resolved liver as the most sensitive organ and reduced interstudy variability: liver BMDs spanned ∼80-fold in daily-dose units but only ∼20-fold when scaled to cumulative dose. Among closely-matched mouse liver gavage studies, cumulative-dose BMDs only varied by ∼2.5-fold across 1 to 28-day treatment regimens. Error-corrected sequencing also demonstrated parity, with acute-dose regimens producing mutation burdens near-identical (< 1.2-fold) to those cumulated from 28-day repeat-dose regimens. Comparable results were obtained from carcinogenicity datasets confirming proportionality-of-effect to cumulative dose. These findings empirically support the validity of the LTL-TTC concept. More broadly, they demonstrate that short-term *in vivo* mutagenicity assays can serve as reliable surrogates for lifetime carcinogenicity studies, strengthening the scientific and regulatory basis for duration-adjusted acceptable intakes for *N-*nitrosamine impurities.

## INTRODUCTION

The management of *N*-nitrosamine impurities represents an ongoing challenge for industry, academic and regulatory scientists affecting drug development and the global drug supply (Kruhlak et al. 2024). *N*-nitrosamines form under a range of conditions, including as by-products of normal, physiological processes (Lynch et al. 2024). Although their potency spectrum is broad (Thomas et al. 2021; Thresher et al. 2020), many are recognised as mutagenic carcinogens in animals and various organisations have categorised certain *N-*nitrosamines as probable human carcinogens (Horne et al. 2023; IARC 1987). In 2018, the detection of the low-molecular weight nitrosamines *N-*nitrosodimethylamine (NDMA) and *N-*nitrosodiethylamine (NDEA) in some sartan-based drug products led to large-scale recalls and renewed focus on *N-*nitrosamine formation, patient exposure and risk assessment (Ruepp et al. 2021; Sörgel et al. 2019). Upon ingestion, small dialkyl nitrosamines undergo cytochrome P450 (CYP2E1) mediated α-hydroxylation, producing formaldehyde and highly reactive diazonium ions capable of alkylating DNA (Haggerty and Holsapple 1990; Yang et al. 1987). Such adducts can induce mutations if unrepaired (Cross and Ponting 2021). While formaldehyde itself is mutagenic in bacterial and mammalian systems, its carcinogenicity is considered route-specific, with the weight-of-evidence suggesting it is carcinogenic in rodents following inhalation but not oral exposure (EMA 2023).

Traditionally, quantitative human cancer risk assessments have relied on linear extrapolation from 24-month rodent carcinogenicity bioassay data (EMA 2018). Typically, dose-response modelling is used to derive the ‘tumorigenic dose 50’ (TD50) defined as the dose estimated to produce a 50% tumour incidence in rodents. Linear low-dose extrapolation is then applied to this estimate to infer the dose associated with a specified excess lifetime cancer risk (*e.g.,* one in a million) (Thomas et al. 2025). This approach is considered conservative (Johnson et al. 2021), and this has driven the development of the ‘Threshold of Toxicological Concern’ (TTC) concept in the pharmaceutical sector (Kroes et al. 2004). The ICH M7 guideline formalises the TTC application to the control of mutagenic impurities in drug substances and products (EMA 2023). It focuses on DNA-reactive (mutagenic) impurities that are positive in the *in vitro* bacterial mutagenicity (Ames) test or the *in vivo* transgenic rodent gene mutation assay. In the absence of additional, *in vivo* information, the proscribed TTC acceptable intake for mutagenic impurities is 1.5 μg/day. This corresponds to a theoretical one in a hundred-thousand excess lifetime cancer risk, justified based on the specific risk-benefit of medicinal use in patients (EMA 2023; Kroes et al. 2004; Snodin 2023).

The original TTC concept was established for daily, lifetime exposures (70 years) which, alongside concepts of controlling impurity levels as described in ICH M7 (R2), form two of the three foundational pillars for the control of mutagenic impurities in pharmaceuticals (EMA 2023). A third pillar addresses exposures that are ‘less-than-lifetime’, encompassing short-term, intermediate, and intermittent dosing regimens (Bercu et al. 2021; EMA 2023). A relationship between exposure duration and toxic potency has long been recognised: since the 1920s, Haber’s law has expressed toxicity as a function of total exposure (*i.e., C* x *t*, where *C* is concentration and *t* is time) (Gaylor 2000; Haber 1924). Building on this, the less-than-lifetime concept has existed in regulatory frameworks since the 1980s when U.S. EPA guidance stated that it can be assumed that a high dose of a carcinogen received over a less-than-lifetime scenario is equivalent to a corresponding low dose spread over a lifetime when the total exposure is equivalent (*i.e., C* x *t* = *k*, where *k* is a constant) (Bercu et al. 2021). Standard cancer risk estimates thus rely on the average lifetime daily dose, which has been shown, both theoretically and empirically, to be valid within a factor of twenty for carcinogenesis (Gaylor 2000). Consistent with this, ICH M7 (R2) states that *“standard risk assessments of known carcinogens assume that cancer risk increases as a function of cumulative dose”* (EMA 2023). On this basis, the guideline introduces a less-than-lifetime TTC approach, defining progressively lower daily acceptable intake limits for longer exposures (< 1-month, 1-month to 1-year, 1-year to 10-years and > 10 years) (EMA 2023). Empirical analyses of *N-*nitrosamine rodent carcinogenicity data using TD50 modelling support this principle, showing that cancer incidence correlates with cumulative exposure duration (Bercu et al. 2021; Felter et al. 2025), and that NDEA satisfies the LTL–TTC framework without exceeding a negligible incremental cancer risk (Bercu et al. 2021).

Within ICH M7, nitrosamines are designated as a “cohort of concern” because of their perceived mutagenic and carcinogenic potency, and thus the guideline recommends compound-specific assessment rather than direct TTC application (EMA 2023). This has driven extensive use of transgenic rodent (TGR) gene-mutation assays, which offer increased 3Rs alignment and a practical alternative to the rodent cancer bioassay given the growing realisation that shorter-term genotoxicity and mode-of-action studies can effectively predict carcinogenic risk in humans (Johnson et al. 2021; Marone et al. 2014). TGR studies thus provide critical evidence of *in vivo* mutagenicity and have demonstrated concordance with the two-year rodent bioassay for genotoxic carcinogens (Lambert et al. 2005; Zeller et al. 2018). Consequently, TGR data now play a pivotal role in the assessment of potentially mutagenic impurities for regulatory purposes (EMA 2023).

Despite this progress, the absence of empirical data for many complex nitrosamines (*e.g., N-*nitrosamine drug-substance related impurities or NDSRIs) poses challenges for establishing robust intake limits which can ultimately restrict patient access to medicines (Johnson et al. 2021; Kruhlak et al. 2024; Snodin 2023). To address this, the ‘Carcinogenic Potency Categorisation Approach’ (CPCA) was developed (Kruhlak et al. 2024). This is a structure-activity based method that assigns nitrosamines to one of five decreasing potency categorisations, each with a corresponding intake limit (Kruhlak et al. 2024). While adopted by several regulatory agencies, the CPCA was primarily designed as a starting point framework (Kruhlak et al. 2024) in absence of compound-specific data. Moreover, current CPCA limits are fixed and do not account for exposure duration, meaning that short-term treatments are subject to the same limits as lifetime therapies.

The TD50 model, while widely used for deriving points of departure (PoDs) from carcinogenicity data, was not designed to account for biological compensatory processes (*e.g.,* DNA repair) that shape the sigmoidal nature of many dose-response relationships (Thomas et al. 2025). In this regard, it is recognised that a more recent approach called benchmark dose (BMD) modelling offers a more flexible and statistically robust alternative (Thomas et al. 2025; Wills et al. 2016a). The BMD approach (**Fig. 1a**) works by fitting models to a dose-response relationship. The ‘benchmark dose’ most likely to elicit some predetermined benchmark response (*e.g.,* a 50% increase in response relative to vehicle control levels) is then estimated from the best-fitting model curve. By also considering other plausible fits to the data (*i.e.,* using different model families and the variability in response between animals in each dose-group) the model-average BMD confidence interval can be established (Wills et al. 2016a). This is described by the ‘BMDL’ and ‘BMDU’ values, which represent the lower and upper confidence limits of the BMD, respectively. In this way, BMD estimates and their BMDL – BMDU confidence intervals represent quantitative potency estimates with uncertainty context that reflect the ‘quality’ of the underlying dose-response data (Wills et al. 2016a; Wills et al. 2016b).

**Fig. 1-.**
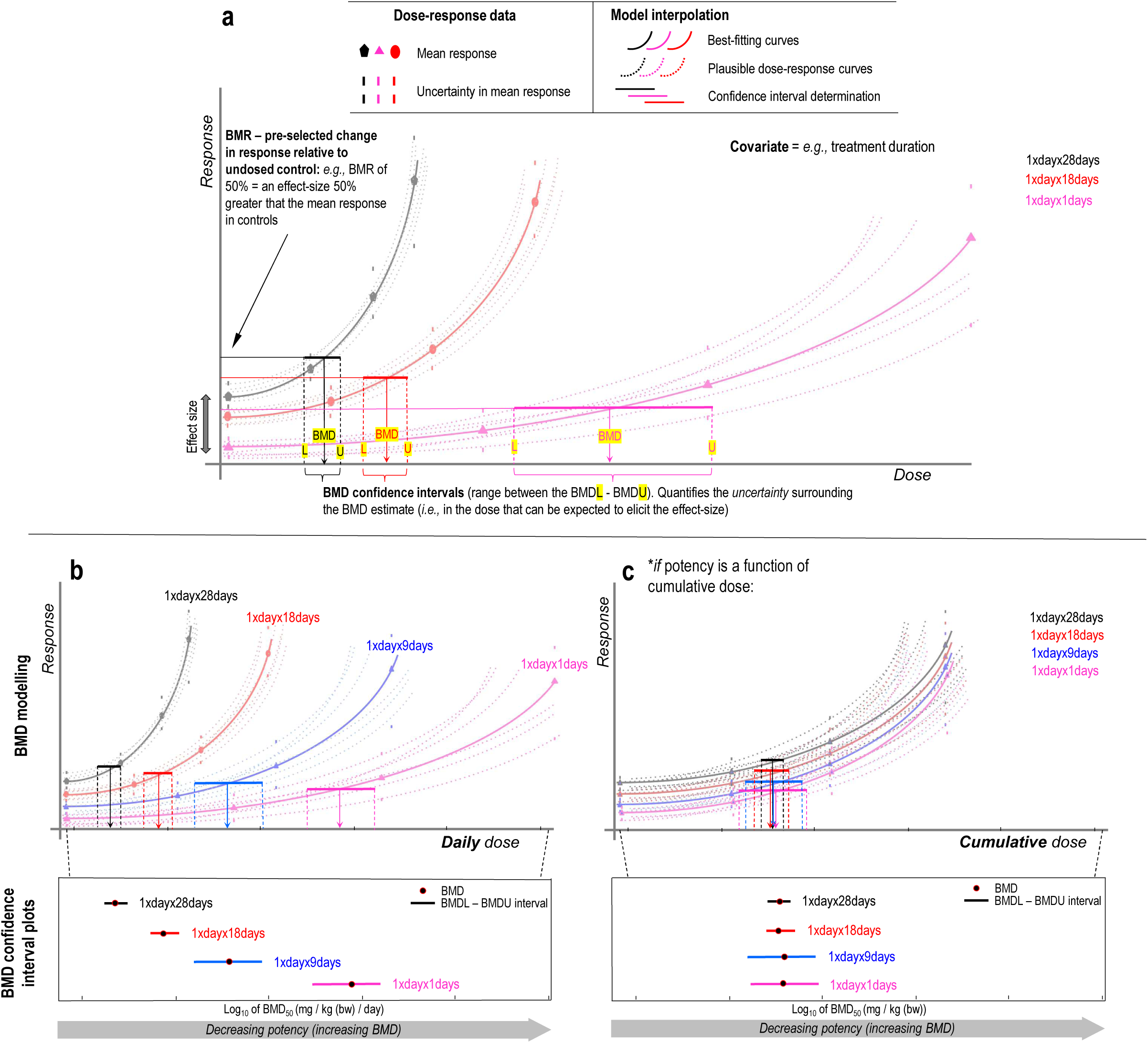
Examining less-than-lifetime (LTL) potency relationships using benchmark dose modelling. **(a)** Benchmark dose (BMD) modelling uses curve-fitting to estimate the dose that can be expected to elicit a pre-specified, small effect-size termed the ‘benchmark response’ (BMR) relative to vehicle control. Whereas the best-fitting model (solid curve) will give the best estimate of the BMD, consideration of other plausible fits to the dose-response data (dashed curves) allows the lower and upper BMD confidence limits to be established (*i.e.,* represented by the BMDL – BMDU interval). An approach termed combined, covariate BMD modelling allows multiple dose-response relationships to be assessed simultaneously using a covariate (*e.g.,* treatment duration) to identify the individual datasets. Because BMD confidence intervals represent estimates of the dose that will elicit a common effect-size, they are quantitative potency estimates and thus well-suited to enable robust potency comparisons within an endpoint. This is demonstrated schematically for the example of different, repeat-dose treatment regimens by the BMD confidence interval plots shown in panels (**b**/**c)**. Panel (**b**) demonstrates the case wherein longer, daily treatment regimens yield more potent responses (*i.e.,* the more days the treatment regimen is repeated over, the lower the BMD estimate in daily-dose units). This example reflects the less-than-lifetime concept, wherein the consequence of longer-term, daily exposures of the same dose is greater than for shorter-term daily exposures. (**c**) By re-expressing the example shown in (**b**) in terms of the total, cumulative dose administered across the entire treatment duration, the ‘equivalence’ of each response to these different exposure durations can be established. The central tenet of the less-than-lifetime threshold of toxicological concern (TTC) – as described in the ICH M7 guida1nce – is that “potency is a function of cumulative dose”. *If* this is the case, the BMD estimates on a cumulative-dose scale would be expected to be approximately equal (**c**) presenting as a vertical stack of similar BMD estimates with overlapping confidence intervals

Work by multiple authors has shown that comparing BMD confidence intervals represents the most robust way to rank and compare potencies and tissue sensitivities within an endpoint (Slob and Setzer 2014; Wills et al. 2016a; Wills et al. 2016b). Whereas the BMD represents the dose-estimate most likely to elicit the benchmark response size, comparing confidence intervals is important because they define the range within which we are most assured that the true BMD lies – based on the uncertainties in the data. A consequence of this is that BMD values arising across datasets should only be considered significantly different when confidence intervals do not overlap. When intervals do overlap, the interpretation is that the underlying dose-response relationships do not contain sufficient information to define how potencies differ (Wills et al. 2016a; Wills et al. 2016b).

In recent years, combined modelling approaches (*e.g.,* the combined-covariate BMD approach) have been developed, enabling simultaneous analysis of multiple dose-response datasets for a shared endpoint using a covariate factor (*e.g.,* treatment duration, sampling time, compound, strain *etc.*) to identify the individual datasets. Importantly, the combined-covariate approach can provide more precise BMD estimates when the shapes of the normalised dose-responses are similar enough at the covariate level to enable parameter estimation (*i.e.,* typically the maximum response ‘*c*’ and log-steepness ‘*d*’ parameters for continuous endpoints) using *all* data in the combined analysis collectively (Slob and Setzer 2014; Wills et al. 2016a; Wills et al. 2016b).

Through this lens, combined BMD analysis provides a powerful method to empirically test the central tenet of the LTL–TTC approach – that cancer risk, and by extension mutagenic potency, scales with cumulative dose (schematically explained, **Fig. 1b/c**). Here, we investigate these less-than-lifetime potency relationships in context of tissue sensitivity, sampling timepoint and route of administration for the data-rich nitrosamine *N-*nitrosodimethylamine (NDMA) using meta-analysis of dose-response data from transgenic rodent, error-corrected sequencing and rodent carcinogenicity studies.

## MATERIALS & METHODS

### Data collection: in vivo mutagenicity assays

Transgenic rodent (TGR) assay dose-response data for NDMA were collected from a GSK study (Lynch et al. 2024) and by literature review. To search the published literature, the search term ((“MutaMouse” OR “Big Blue” OR “lacZ” OR “lacI” OR “gpt” OR “cII”) AND (mutation OR mutant OR mutagen OR mutagenesis) AND (mouse OR rat OR rodent OR mice OR rats OR rodents) AND (NDMA OR DMN OR nitrosodimethylamine OR dimethylnitrosamine)) was used. Identified studies (cut-off date August 2025) were then screened manually to identify dose-response datasets suitable for BMD modelling (*i.e.,* containing at least two dose-groups plus control). These TGR datasets and their sources (Butterworth et al. 1998; Gollapudi et al. 1998; Hobbie et al. 2009; Jiao et al. 1997; Lynch et al. 2024; Mirsalis et al. 1993; Souliotis et al. 1998; Suzuki et al. 1996) are summarised in **Table 1**. Experimentally-matched (*i.e.,* using tissues from the GSK TGR study (Lynch et al. 2024)) error-corrected sequencing data were collected from Ashford *et al.,* 2025 (Ashford et al. 2025) using TwinStrand Biosciences’s duplex sequencing method.

**Table 1.** NDMA transgenic rodent studies with data suitable for benchmark dose modelling.

| Study No | Species | TGR | Trans-gene | N per group | Dose-groups (daily units) mg/kg(bw)/day | Dose-groups (cumulative units) mg/kg(bw) | No. dose-groups | Exposure.days | Sampling.day | Tissue | Sex | Age (weeks) | Route | Data Type | Regimen | Source |
| --- | --- | --- | --- | --- | --- | --- | --- | --- | --- | --- | --- | --- | --- | --- | --- | --- |
| 1 | mouse | BigBlueMouse | lacI | 10 | 0/2/4/8 | 0/8/16/32 | 3.dose.plus.control | 4 | 10 | Liver | f | 9 | gav | sum | 1dayx4days | Butterworth *et al.*, 1998 |
| 2 | mouse | BigBlueMouse | lacI | 10 | 0/2/4 | 0/8/16 | 2.dose.plus.control | 4 | 30 | Liver | f | 9 | gav | sum | 1dayx4days | Butterworth *et al.*, 1998 |
| 3 | mouse | BigBlueMouse | lacI | 10 | 0/2/4 | 0/8/16 | 2.dose.plus.control | 4 | 90 | Liver | f | 9 | gav | sum | 1dayx4days | Butterworth *et al.*, 1998 |
| 4 | mouse | BigBlueMouse | lacI | 10 | 0/2/4 | 0/8/16 | 2.dose.plus.control | 4 | 180 | Liver | f | 9 | gav | sum | 1dayx4days | Butterworth *et al.*, 1998 |
| 5 | rat | BigBlueRat | lacI | 4 | 0/0.2/0.6/2/6 | 0/1.8/5.4/18/54 | 4.dose.plus.control | 9 | 12 | Liver | m | 12 | gav | idv | 1dayx9days | Gollapudi *et al.*, 1998 |
| 6 | rat | BigBlueRat | cII | 4 | 0/0.2/0.6/2/6 | 0/1.8/5.4/18/54 | 4.dose.plus.control | 9 | 12 | Liver | m | 21 | gav | idv | 1dayx9days | Gollapudi *et al.*, 1998 |
| 7 | mouse | MutaMouse | lacZ | 5 | 0/2/6/10 | 0/2/6/10 | 3.dose.plus.control | 1 | 30 | BoneMarrow | m | 15 | gav | sum | 1dayx1days | Jiao *et al.*, 1997 |
| 8 | mouse | MutaMouse | lacZ | 5 | 0/2/6/10 | 0/2/6/10 | 3.dose.plus.control | 1 | 90 | BoneMarrow | m | 15 | gav | sum | 1dayx1days | Jiao *et al.*, 1998 |
| 9 | mouse | MutaMouse | lacZ | 5 | 0/2/6/10 | 0/2/6/10 | 3.dose.plus.control | 1 | 30 | Liver | m | 15 | gav | sum | 1dayx1days | Jiao *et al.*, 1998 |
| 10 | mouse | MutaMouse | lacZ | 5 | 0/2/6/10 | 0/2/6/10 | 3.dose.plus.control | 1 | 90 | Liver | m | 15 | gav | sum | 1dayx1days | Jiao *et al.*, 1998 |
| 11 | mouse | BigBlueMouse | lacI | 4 | 0/4/6 | 0/20/30 | 2.dose.plus.control | 5 | 14 | Liver | m | 3 | ip | idv | 1dayx5days | Mirasalis *et al.*, 1993 |
| 12 | mouse | BigBlueMouse | lacI | 4 | 0/4/6 | 0/20/30 | 2.dose.plus.control | 5 | 14 | Liver | m | 6 | ip | idv | 1dayx5days | Mirasalis *et al.*, 1993 |
| 13 | mouse | BigBlueMouse | lacI | 4 | 0/4/6 | 0/20/30 | 2.dose.plus.control | 5 | 14 | Liver | m | 3 | ip | idv | 1dayx5days | Mirasalis *et al.*, 1993 |
| 14 | mouse | BigBlueMouse | lacI | 4 | 0/4/6 | 0/20/30 | 2.dose.plus.control | 5 | 14 | Liver | m | 6 | ip | idv | 1dayx5days | Mirasalis *et al.*, 1993 |
| 15 | mouse | MutaMouse | lacZ | 6 | 0/0.02/0.07/0.19/0.36/1.1/2/4 | 0/0.56/1.96/5.32/10.08/30.8/56/112 | 6.dose.plus.control | 28 | 31 | Liver | m | 8-11 | gav | idv | 1dayx28days | Lynch *et al.*, 2024 |
| 16 | mouse | MutaMouse | lacZ | 6 | 0/5/10 | 0/5/10 | 2.dose.plus.control | 1 | 31 | Liver | m | 8-11 | gav | idv | 1dayx1days | Lynch *et al.*, 2024 |
| 17 | mouse | MutaMouse | lacZ | 5 | 0/2/4 | 0/56/112 | 2.dose.plus.control | 28 | 31 | Bladder | m | 8-11 | gav | idv | 1dayx28days | Lynch *et al.*, 2024 |
| 18 | mouse | MutaMouse | lacZ | 3-5 | 0/2/4 | 0/56/112 | 2.dose.plus.control | 28 | 31 | Stomach | m | 8-11 | gav | idv | 1dayx28days | Lynch *et al.*, 2024 |
| 19 | mouse | MutaMouse | lacZ | 3-6 | 0/0.36/1.1/2/4 | 0/10.08/30.8/56/112 | 4.dose.plus.control | 28 | 31 | Kidney | m | 8-11 | gav | idv | 1dayx28days | Lynch *et al.*, 2024 |
| 20 | mouse | MutaMouse | lacZ | 3-6 | 0/0.36/1.1/2/4 | 0/10.08/30.8/56/112 | 4.dose.plus.control | 28 | 31 | Lung | m | 8-11 | gav | idv | 1dayx28days | Lynch *et al.*, 2024 |
| 21 | mouse | MutaMouse | lacZ | 4 | 0/0/5/10 | 0/0/5/10 | 2.dose.plus.2.controls | 1 | 28 | Liver | m | 9.5 | ip | sum | 1dayx1days | Souliotis *et al.*, 1998 |
| 22 | mouse | MutaMouse | lacZ | 4 | 0/0/5/10 | 0/0/5/10 | 2.dose.plus.2.controls | 1 | 28 | BoneMarrow | m | 9.5 | ip | sum | 1dayx1days | Souliotis *et al.*, 1998 |
| 23 | mouse | MutaMouse | lacZ | 4 | 0/0/5/10 | 0/0/5/10 | 2.dose.plus.2.controls | 1 | 14 | Liver | m | 9.5 | ip | sum | 1dayx1days | Souliotis *et al.*, 1998 |
| 24 | mouse | MutaMouse | lacZ | 4 | 0/0/5/10 | 0/0/5/10 | 2.dose.plus.2.controls | 1 | 14 | BoneMarrow | m | 9.5 | ip | sum | 1dayx1days | Souliotis *et al.*, 1998 |
| 25 | mouse | MutaMouse | lacZ | 4 | 0/0/5/10 | 0/0/5/10 | 2.dose.plus.2.controls | 1 | 28 | Lung | m | 9.5 | ip | sum | 1dayx1days | Souliotis *et al.*, 1998 |
| 26 | mouse | MutaMouse | lacZ | 4 | 0/0/5/10 | 0/0/5/10 | 2.dose.plus.2.controls | 1 | 14 | Lung | m | 9.5 | ip | sum | 1dayx1days | Souliotis *et al.*, 1998 |
| 27 | mouse | MutaMouse | lacZ | 4 | 0/0/5/10 | 0/0/5/10 | 2.dose.plus.2.controls | 1 | 28 | Spleen | m | 9.5 | ip | sum | 1dayx1days | Souliotis *et al.*, 1998 |
| 28 | mouse | MutaMouse | lacZ | 4 | 0/0/5/10 | 0/0/5/10 | 2.dose.plus.2.controls | 1 | 14 | Spleen | m | 9.5 | ip | sum | 1dayx1days | Souliotis *et al.*, 1998 |
| 29 | mouse | BigBlueMouse | lacI | 3 | 0/5/10 | 0/5/10 | 2.dose.plus.2.controls | 1 | 14 | Liver | m | 7 | ip | idv | 1dayx1days | Suzuki *et al.*, 1996 |
| 30 | rat | BigBlueRat | cII | 4 | 0/0.005/0.05/0.25/0.5/1.26 | 0/0.07/0.7/3.5/7/17.64 | 5.dose.plus.control | 14 | 30 | Liver | m | 4 | drink | idv | drink.14days | Hobbie *et al.*, 2009 |

### Data collection: rodent cancer bioassay

Rodent cancer bioassay data for NDMA were identified from the Agency for Toxic Substances and Disease Registry’s (ATSDR) Toxicological Profile for *N-*Nitrosodimethylamine (ATSDR 2023), the Carcinogenic Potency Database (CPDB), alongside additional searches of the published literature (CPDB 2011). Rodent carcinogenicity studies concluded positive by the CPDB or where the authors concluded the tumour incidences were the result of a treatment related effect were considered. Studies were required to meet the original CPDB study inclusion criteria, however, studies of any exposure duration were considered if the overall experimental duration was at least a quarter of the standard lifespan (*i.e.,* inclusion of rodent studies where the overall study duration was ≥ 26 weeks) and only studies where the effective number of animals per group was ≥ 10 were selected for further analysis (Bercu et al. 2021). The most sensitive organ site was identified for each species and sex. The total tumour incidence for each organ site (totalling all lesions including adenomas and carcinomas) was collected for analysis. Where dose was not reported in mg/kg(bw)/day it was converted using CPDB standard values for dose-calculation (CPDB). Rodent carcinogenicity dose-response datasets and their sources (Engelse et al. 1974; Keefer et al. 1973; Lijinsky and Reuber 1984; Peto et al. 1991b; Takahashi et al. 2000; Tanaka et al. 1988; Terracini et al. 1967) are summarised in **Table 2**.

**Table 2.** NDMA rodent-cancer bioassay studies with data suitable for benchmark dose modelling.

| Study No. | Species | Strain | N per group | Dose-groups (daily units) mg/kg(bw)/day | Dose-groups (cumulative units) mg/kg(bw) | No. dose-groups | Exposure weeks | Sampling weeks | Tumor site | Sex | Route | Data Type | Regimen | Source |
| --- | --- | --- | --- | --- | --- | --- | --- | --- | --- | --- | --- | --- | --- | --- |
| 1 | rat | F344 | 20 | 0 / 0.314 / 0.743 | 0 / 47.1 / 111.45 | 2.dose.plus.control | 30 | lifetime | Liver | f | Water | quantal | 5xweekx30weeks | Lijinsky et al., 1984 |
| 2 | rat | Wistar | 11 | 0 / 0.5 / 1.25 | 0 / 105 / 262.5 | 2.dose.plus.control | 30 | 30 | Liver | m | Water | quantal | 1xdayx30weeks | Takahashi et al., 2000 |
| 3 | mouse | C3Hf | 20 | 0 / 1.24 / 1.35 | 0 / 112.84 / 122.85 | 2.dose.plus.control | 13 | 58 | Liver | f | Water | quantal | 1xdayx13weeks | DenEngelse et al., 1974 |
| 4 | rat | MRC | 15 or 30 | 0 / 0.2 / 1 | 0 / 30 / 150 | 2.dose.plus.control | 30 | 100 | Liver | m | Water | quantal | 5xweekx30weeks | Keefer et al., 1973 |
| 5 | rat | Colworth | 240 or 60 | 0 / 0.001 / 0.003 / 0.005 / 0.011 / 0.022 / 0.044 / 0.065 / 0.087 / 0.109 / 0.131 / 0.174 / 0.218 / 0.261 / 0.348 / 0.697 | 0 / 0.73 / 2.18 / 3.64 / 8.01 / 16.02 / 32.03 / 47.32 / 63.34 / 79.35 / 95.37 / 126.67 / 158.7 / 190.01 / 253.34 / 507.42 | 15.dose.plus.control | lifetime | lifetime | Liver | m | Water | quantal | 1xdayxlifetime | Peto et al., 1991 |
| 6 | rat | Colworth | 240 or 60 | 0 / 0.002 / 0.005 / 0.01 / 0.019 / 0.038 / 0.076 / 0.115 / 0.153 / 0.191 / 0.229 / 0.306 / 0.382 / 0.459 / 0.612 / 1.224 | 0 / 1.46 / 3.64 / 7.28 / 13.83 / 27.66 / 55.33 / 83.72 / 111.38 / 139.05 / 166.71 / 222.77 / 278.1 / 334.15 / 445.54 / 891.07 | 15.dose.plus.control | lifetime | lifetime | Liver | f | Water | quantal | 1xdayxlifetime | Peto et al., 1991 |
| 7 | rat | Porton | 6 - 19 | 0 / 0.08 / 0.2 | 0 / 58.24 / 145.6 | 2.dose.plus.control | lifetime | 104-120wks | Liver | m | Diet | quantal | 1xdayxlifetime | Terracini et al., 1967 |
| 8 | rat | Porton | 5 - 62 | 0 / 0.1 / 0.25 / 0.5 / 1 / 2.5 | 0 / 72.8 / 182 / 364 / 728 / 1820 | 5.dose.plus.control | lifetime | 104-120wks | Liver | f | Diet | quantal | 1xdayxlifetime | Terracini et al., 1967 |
| 9 | hamster | Syrian | 16 - 26 | 0 / 6 / 12 | 0 / 90 / 180 | 2.dose.plus.control | 15 | lifetime | Liver | m | Intratracheal | quantal | 1xweekx15weeks | Tanaka et al., 1988 |

### Benchmark dose analyses

Benchmark dose analyses were carried out using the PROAST R-package (version 70.9) in the R programming environment (version 4.1.1) (RIVM 2025). As established in previous work, for both TGR and rodent carcinogenicity outputs, confidence interval plots ordered by BMD were employed to visually compare potencies across different studies whilst taking estimation uncertainty into account (Wills et al. 2016b).

### Continuous BMD analyses: transgenic rodent data

Unless stated otherwise, data were analysed using the four-model average method using the ‘canonical’ model families (*i.e.,* exponential, Hill, inverse exponential and log-normal models) suitable for relative potency comparisons (Slob et al. 2025). Combined analyses (*i.e.,* analyses of multiple dose-response relationships simultaneously) were conducted by exposure route and tissue utilising ‘substudy’ as the covariate factor. To prevent overfitting, more complex models with additional parameters were only accepted if the goodness-of-fit exceeded the critical value at P < 0.05. Model-average bootstrap sampling used 200 simulations per subgroup. PROAST outputs designate potency (*i.e.,* the BMD) and its two-sided 90% confidence interval (*i.e.,* the BMDL and BMDU) as the ‘critical effect dose’ (*i.e.,* CED, CEDL and CEDU), respectively, for each level of the covariate. Based on detailed analyses (White et al. 2025) and expert working-group recommendations the benchmark response (BMR) (*i.e.,* the critical effect-size or CES) used in the presented analyses was 50% (*i.e.,* denoting a 50% increase in mutant frequency response relative to concurrent control) (EFSA 2022). The CEDL and CEDU values represent the lower and upper bounds of the two-sided 90% confidence interval of the BMD, respectively, with the difference between the CEDU and the CEDL defining the ‘width’ of the confidence interval, and therefore, its precision.

### Quantal BMD analyses: rodent cancer bioassay data

Data were analysed using the PROAST quantal ‘model-average’ (exponential, Hill, two-stage, log-logistic, Weibull, log-probit and gamma) workflow. Combined analyses (*i.e.,* analyses of multiple dose-response relationships simultaneously) utilised ‘substudy’ as covariate. To prevent overfitting, more complex models with additional parameters were only accepted if the goodness-of-fit exceeded the critical value at P < 0.05. PROAST outputs from quantal data designate potency and its two-sided 90% confidence interval as the ‘Benchmark dose’ (*i.e.,* BMD, BMDL and BMDU), respectively, for each level of the covariate. Based on current recommendations (EFSA 2022), the benchmark response used in the presented analyses was 10% extra risk, defined as a 10% increase in liver tumour incidence relative to background, scaled to the fraction of animals not exhibiting tumours in the control group (*i.e.,* (*P*(*d*) – *P*(0)) / [1 – *P*(0)]). The BMDL and BMDU values represent the lower and upper bounds of the two-sided, 90% confidence interval of the BMD, respectively, with the difference between the BMDU and the BMDL defining the ‘width’ of the confidence interval, and therefore, its precision.

## RESULTS

### BMD-derived tissue sensitivity ranking using the GSK MutaMouse study

In 2024, GSK published the results of a TGR study of NDMA using MutaMouse (Lynch et al. 2024). Statistically significant dose-response relationships were observed for mutant frequency responses in liver, lung and kidney tissues whereas bladder, bone marrow, spleen and stomach were negative (28-day repeat-dose study, 0 – 4 mg/kg(bw)/day). **Fig. 2a** shows BMD_50_ analyses for the three positive tissues. Comparison of the BMD confidence intervals shows the dose required to increase the mutant frequency response by 50% from vehicle control levels in the liver was ∼2 – 9 times lower than in the kidney or lung. Because the liver confidence interval does not overlap with those defined for lung or kidney, we can conclude that this heighted sensitivity is significant. When carrying out this BMD analysis, the datasets for each tissue were analysed one-at-a-time, sequentially, and not using ‘tissue’ as a covariate in a combined analysis. This is because these data are not truly independent of one another – as multiple tissue samples were collected from each animal. When this occurs, it is likely that the response data arising from tissues originating from the same animal will be more closely related (*e.g.,* due to within-animal exposure correlation). Ignoring this can result in inaccurate or overly-optimistic confidence intervals hence the use of the independent, individual-analysis approach in this specific instance (Long et al. 2018). The benchmark dose analyses underlying **Fig. 2a** are presented in **Supplementary Fig. 1**.

**Fig. 2-.**
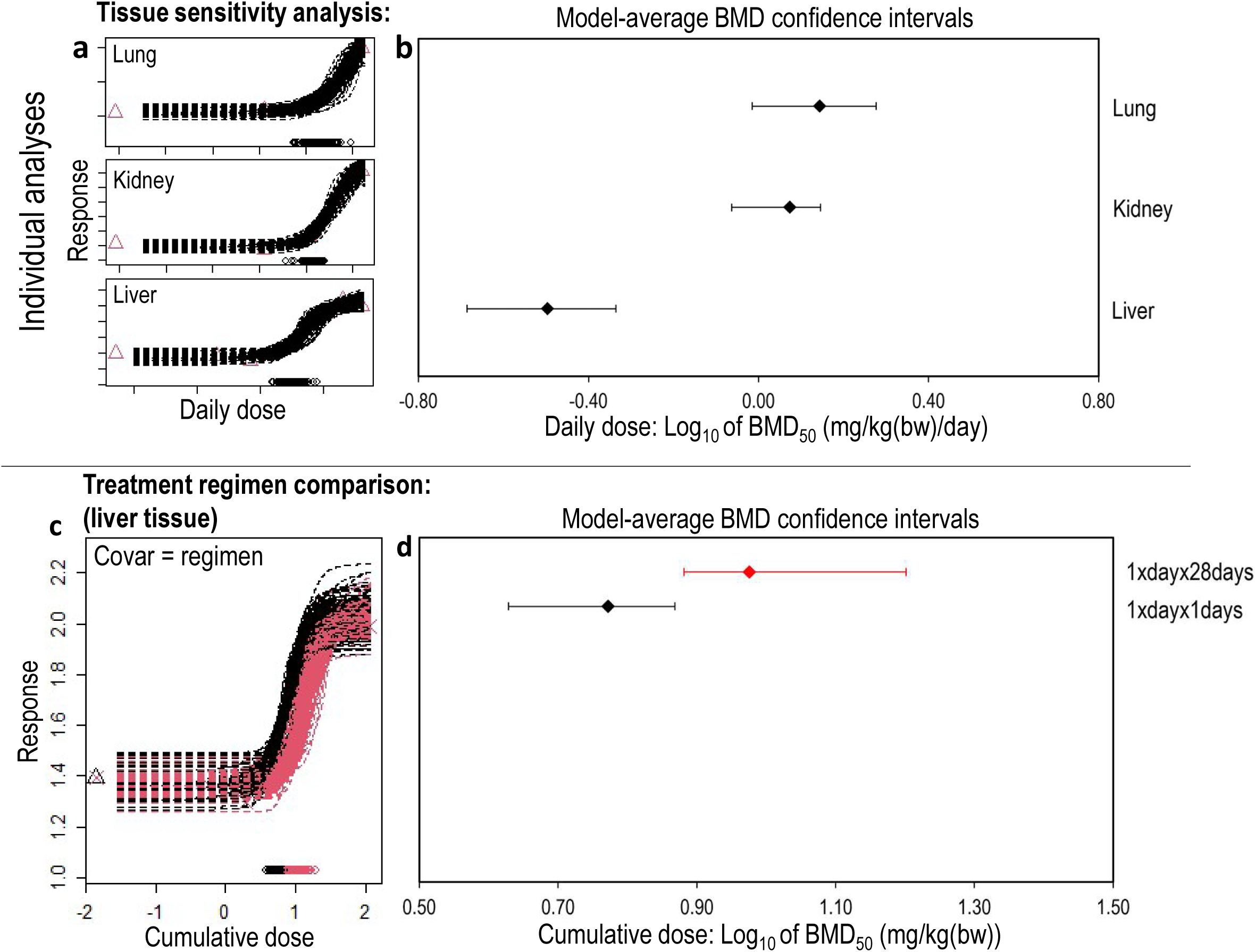
BMD-derived potency comparisons for NDMA using TGR data from the GSK MutaMouse study (Lynch *et al.,* 2024). (**a)** Independent BMD analyses of the dose-response data arising in each of the three positive tissues establish liver as the most sensitive to NDMA-induced mutagenesis (*i.e.,* lowest BMD). (**b)** In liver tissue, combined-covariate BMD analysis on cumulative-dose scale using treatment regimen as a covariate suggests a slight increase in potency when the dose is administered as a single-dose (1xdayx1days) compared to administration using a 28-day, repeat-dose regimen (1xdayx28days)

### Potency comparison of chronic and acute dosing regimens

In the GSK TGR study, the potential to compare mutant frequency responses arising from single-dose or 28-day repeat-dose regimens was incorporated into the study design. Two single administration dose-groups (*i.e.,* 5 or 10 mg/kg; administered once on Day 1) were chosen because they were near-identical to the cumulative doses of two of the 28-day, repeat-dose treatments groups (*i.e.,* 0.19 mg/kg(bw)/day x 28 days’ administration = 5.32 mg/kg and 0.36 mg/kg(bw)/day x 28 days = 10.08 mg/kg) (Lynch et al. 2024). This data provides an opportunity to empirically test the less-than-lifetime concept by comparing the potency of the 28-day, repeat-dose regimen to that of the single-dose regimen on a cumulative dose-scale using treatment regimen as a covariate (**Fig. 2b**). The resultant BMD confidence intervals lie closely together and nearly overlap, indicating similar potencies (and supporting the less-than-lifetime concept of potency as a function of cumulative dose). Comparison of the confidence intervals suggests that the single-dose regimen was just ∼ 1.6 (1.03 – 3.74) times more potent than the cumulated consequence of the 28-day, repeat-dose regimen. The benchmark dose analyses underlying **Fig. 2b** are presented in **Supplementary Fig. 2**.

### Incorporating information from other transgenic rodent studies

To place these findings into broader context, a literature review to identify all NDMA TGR studies with study designs suitable for combined benchmark dose modelling (*i.e.,* two dose plus control or greater) was conducted. The results are summarised in **Table 1**. Including the GSK data analysed above, 30 dose-response datasets arising in seven tissues across three transgenic rodent models, three exposure routes and six treatment regimens were identified.

### Sampling time has no significant effect on BMD

A recognised challenge when comparing TGR-derived dose-response relationships stems from variability in experimental method (*e.g.,* use of different sampling times, exposure routes, species, strains / transgene reporters) across different labs (Wills et al. 2017). Previous work indicates that TGR variant (*i.e.,* assessments using MutaMouse versus Big Blue or gpt delta *etc.*) has no significant effect on the derived BMD, enabling meaningful potency comparisons to be made across TGR variants (Wills et al. 2017). However, the **Table 1** summary shows wide variability in the sampling times utilised (10 – 180 days). The consequence of this in terms of BMD-derived potency estimate is less clearly established by previous studies: if the TGR dose-response is strongly affected by sampling time, it would invalidate meaningful, inter-study potency comparisons across much of the available data.

Fortunately, sampling time was the specific focus of previous TGR investigations (Butterworth et al. 1998) with NDMA wherein the effects of short-term, repeat-dose treatments (2, 4, or 8 mg/kg(bw)/day for four days) followed by sampling on Days 10, 30, 90, or 180 were investigated in female B6C3F_1_ mice (*lacI* locus). These data provide the opportunity to establish the impact of sampling time on BMD-derived potency estimations for NDMA in liver using otherwise exactly-matched experimental conditions. Using a combined analysis with ‘sampling time’ as covariate demonstrated highly similar BMD estimates with overlapping confidence intervals and no obvious temporal trend (**Fig. 3**). This suggests that sampling time can reasonably be ignored as an influential covariate, enabling deeper cross-study potency comparisons of the datasets established from the literature review. From a toxicological standpoint, the analysis also indicates limited contributions to the *lacI* mutant frequency response by selective loss or clonal expansion and identifies that NDMA-induced mutations in the rodent liver persist for extended periods (*i.e.,* ∼25% of mouse life-span) in a manner consistent with slow tissue turn-over and the long half-life of hepatocytes in this species (Long et al. 2018). The benchmark dose analyses underlying **Fig. 3** are presented in **Supplementary Fig. 3**.

**Fig. 3-.**
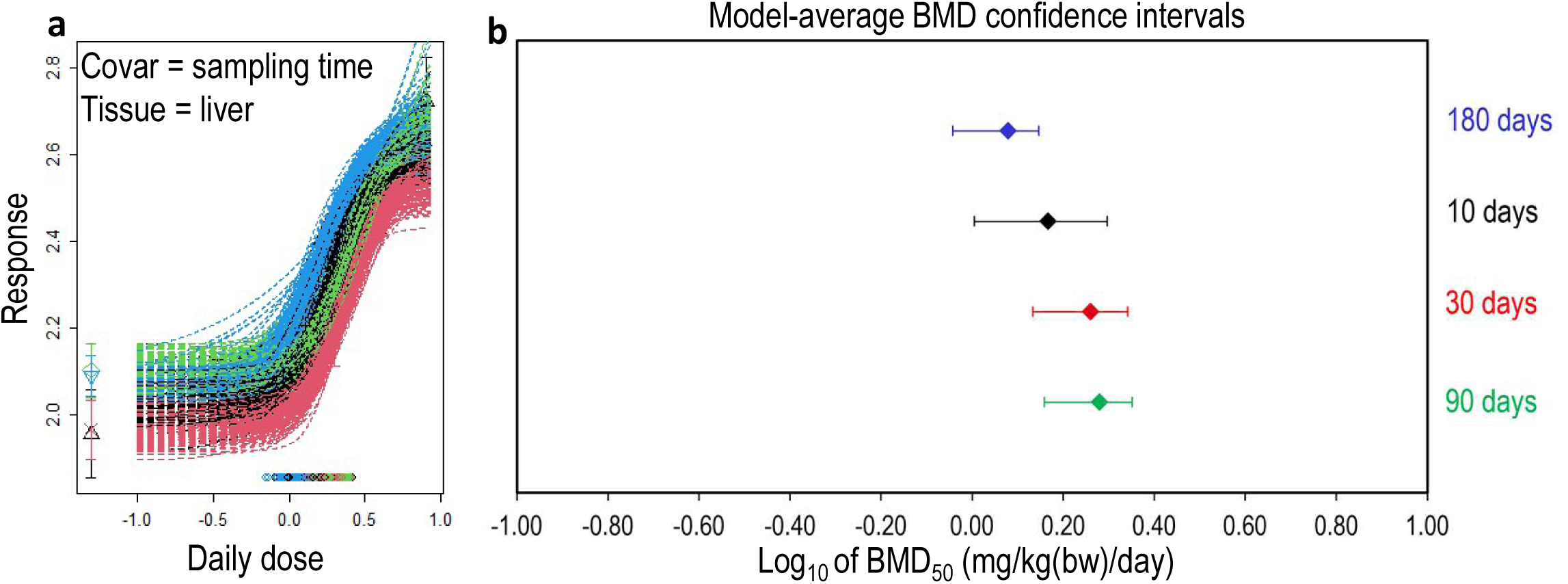
Effect of sampling time on BMD-derived potency estimates using TGR dose-response data for NDMA. **(a)** Combined BMD analysis of four NDMA datasets using ‘sampling time’ as covariate. All datasets used a matched experiment design with a 1xdayx4days treatment regimen, but tissue sampling was carried out on either day 10, 30, 90 or 180. (**b)** BMD confidence interval plot confirming highly similar potency estimates regardless of sampling time with no obvious temporal trend

### BMD analysis of all available TGR data

Having ruled out sampling time as an influential covariate, **Fig. 4** displays the BMD confidence intervals for all identified datasets in **Table 1**, using combined analyses for each exposure route and tissue. With daily dose units on the x-axis, the eighteen BMD estimates from liver tissue span ∼1.9 log units (∼80-fold range on original scales) (**Fig. 4a**). Of course, this wide potency range largely reflects that different treatment durations are not accounted for when dose is described in daily units. Plotting the liver confidence intervals also showed that the only study using drinking water as the administration route exhibited a markedly left-shifted BMD (*i.e.,* a more potent response) compared to the results obtained after gavage or intraperitoneal injection (**Fig. 4a**).

**Fig. 4-.**
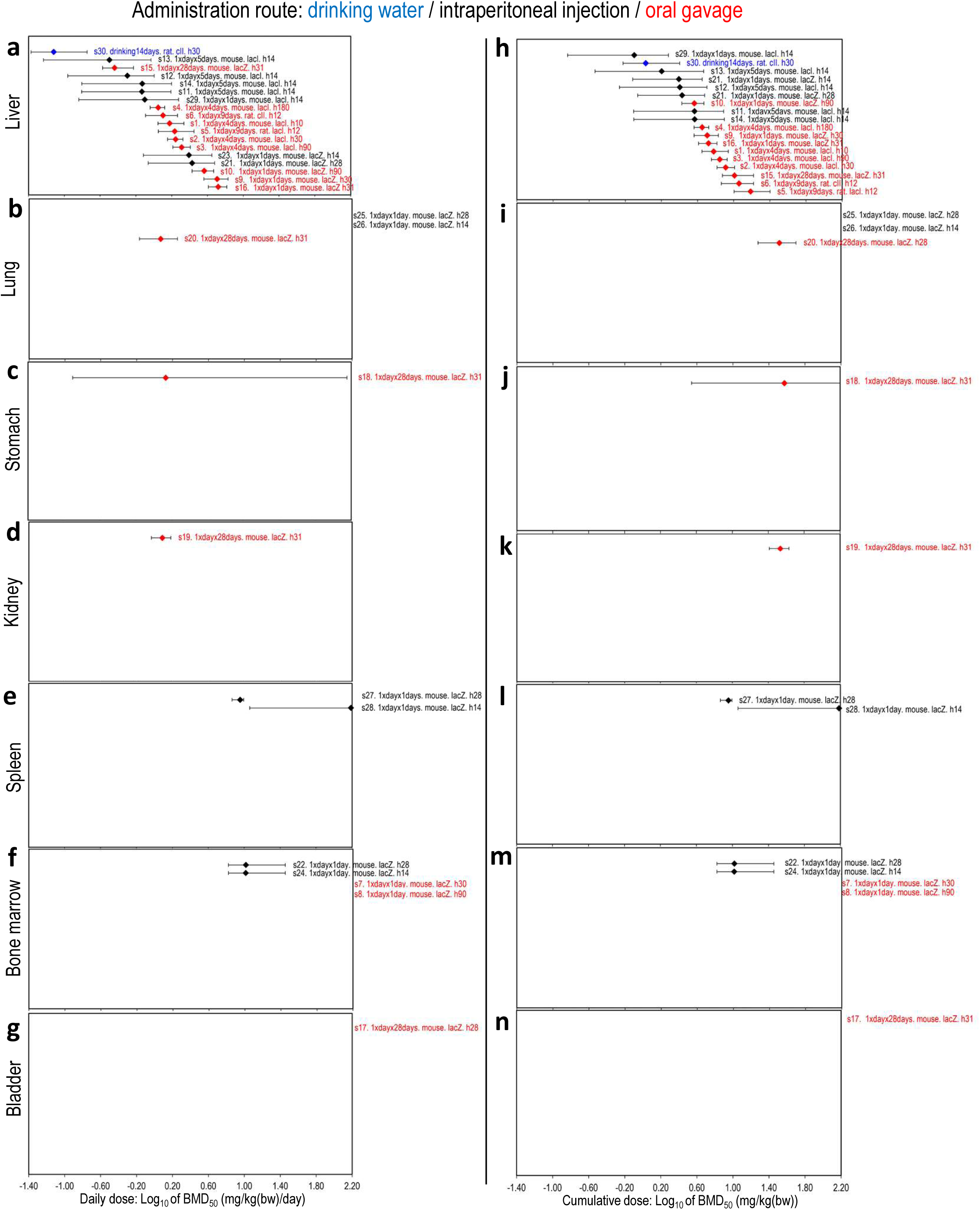
BMD-derived potency estimations for all available TGR data for NDMA. Combined BMD analyses using the exponential model family were carried out for each administration route and tissue with ‘study’ as covariate. (**a-g)** BMD-derived confidence intervals for each tissue on daily-dose scales. (**h-n)** BMD confidence intervals in terms of the total, cumulative dose administered. Confidence interval cut-off by the right-plot boundary indicate datasets for which the upper confidence limit (*i.e.,* the BMDU) was incalculable. Datasets with no visible confidence interval indicate instances where both the upper and lower confidence limits (*i.e.,* the BMDL and BMDU) were incalculable. For each confidence interval, the text indicates the study number (relative to **Table 1**), regimen, species, transgene and tissue harvest time-point used

Looking beyond liver to the results determined in the other tissues (**Fig. 4b-g**), bounded BMDs were calculable for datasets arising in lung, stomach and kidney – albeit with a wide confidence interval in stomach (∼ 3 log units or 1000-fold) indicating considerable uncertainty in the data for this tissue (**Fig. 4c**). In lung, spleen and bone marrow, bounded BMDs were calculable from some datasets whereas others yielded partially-bounded or unbounded BMD estimates (*i.e.,* incalculable BMDUs or incalculable BMDLs and BMDUs). In lung (**Fig. 4b**), single-dose regimens by the intraperitoneal route failed to elicit a dose-response, whereas a bounded BMD was calculable after a 28-day repeat-dose regimen by gavage. In spleen (**Fig. 4e**), the two available datasets originate from the same laboratory, with the only study-design difference being the sampling timepoint. Single doses of 5 or 10 mg/kg were administered by intraperitoneal injection, but sampling at day 14 yielded an unbounded BMDU (*i.e.,* suggesting a weak response at most) whereas sampling at day 28 enabled determination of a fully-bounded BMD. In bone marrow (**Fig. 4b/f**), the results suggest a potential difference due to exposure route. Despite similar, single-dose study designs, via intraperitoneal injection, doses of 5 or 10 mg/kg yielded bounded BMD values at both 14- or 28-day sampling times. In contrast, by gavage dosing, single doses of 2, 6 or 10 mg/kg showed no measurable response yielding an unbounded BMDs at both 30- or 90-day sampling times. Finally, one dataset arising from the GSK study was available for bladder (**Fig. 4g**). Despite repeat-dosing by gavage for 28 days at doses up to 4 mg/kg(bw)/day, no significant dose-response was established.

To better account for the differences in treatment duration, **Fig. 4h-n** explores the potency rankings using cumulative dose-scales. Immediately, liver is more readily established as the most sensitive tissue, and the potent response observed after drinking water administration now falls amongst the other results, suggesting the use of a relatively high-dose regimen for the treatment period (**Fig. 4h**). The upper (*i.e.,* most sensitive) portion of liver potency ranking largely consists of the intraperitoneal studies suggesting the potential for more potent responses via this administration route albeit with considerable confidence interval overlap with many of the gavage outcomes (**Fig. 4h**). The benchmark dose analyses underlying **Fig. 4** are presented in **Supplementary Figs. 4-7.**

### Focused analyses using the liver gavage TGR data

The number of TGR datasets available in liver tissue via the gavage administration route across different daily-treatment durations provides the opportunity to investigate less-than-lifetime potency relationships more closely. **Fig. 5a/b** lays out the model-average BMD confidence intervals on daily-dose scales in order of increasing treatment duration (*i.e.,* in the order 1, 4, 9, then 28 treatment days). The BMD estimates span 1.2 log-units (16-fold on original scales) and, in-line with less-than-lifetime concepts, the BMDs generally decrease with increasing numbers of treatment days (*i.e.,* more potent responses as treatment duration increases – as expected). A slight deviation from this potency trend was observed for the nine treatment-day studies. This might be attributable to species, as these two datasets were collected in rats whereas all other studies were carried out in mice. Switching to cumulative dose-scales (**Fig. 5c/d**), all of the BMD confidence intervals stacked nearly on top of one another, occupying a much narrower range (four-fold across all studies, or just two-fold considering just the eight studies collected in mouse). Collectively, the BMD analyses suggest that cumulative doses of NDMA in the range ∼ 4 – 16 mg/kg(bodyweight) can be expected to increase liver mutant frequency responses by 50% from control-group levels in rodents after daily treatment by gavage for durations between 1- and 28-days.

**Fig. 5-.**
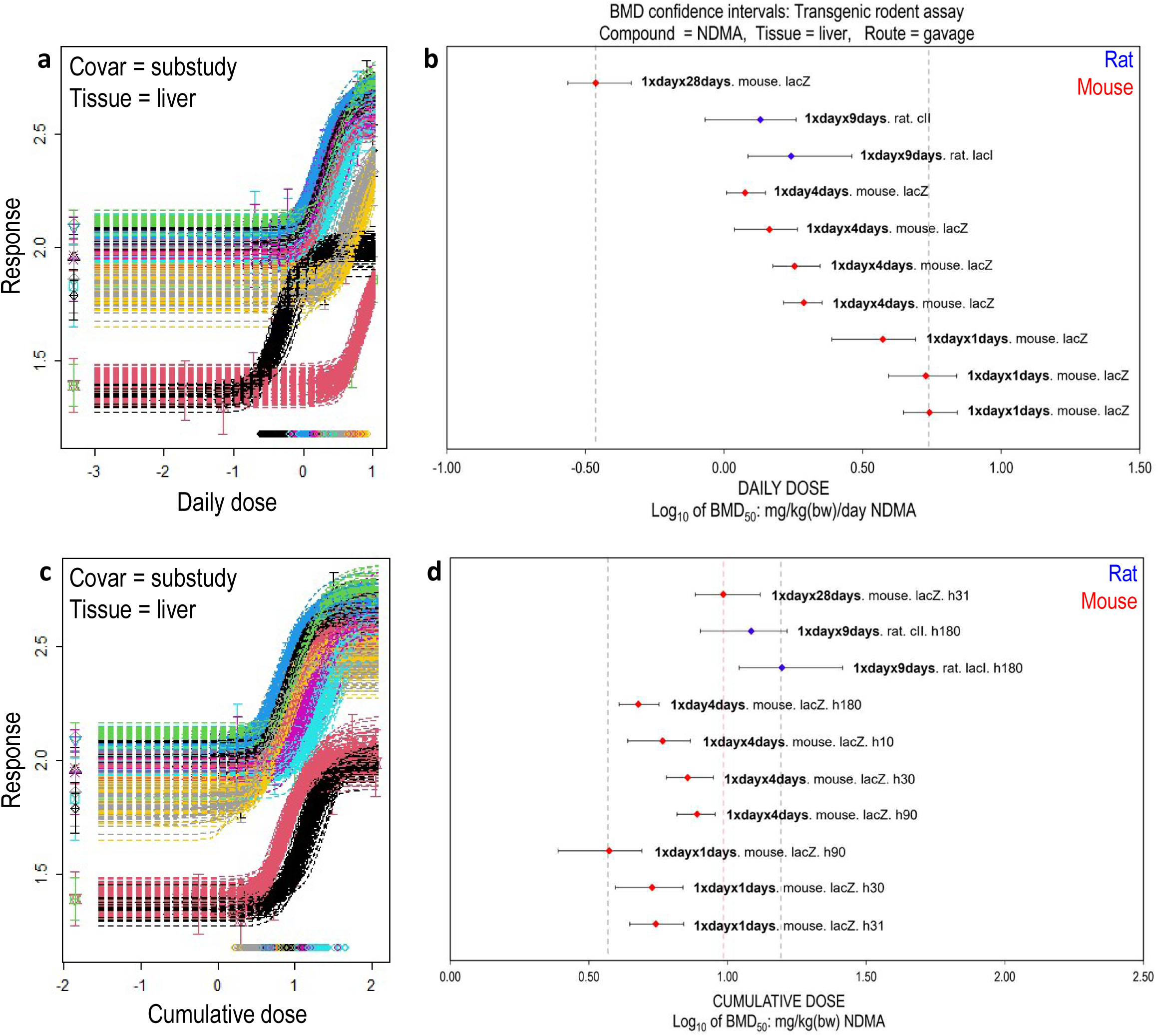
Examining less-than-lifetime potency relationships using all liver TGR data dosed by gavage. **(a)** Model-average BMD analysis and (**b**) confidence intervals on daily-dose scales ranked by increasing treatment duration. In-line with less-than-lifetime concepts, BMDs generally decreased (*i.e.,* demonstrating increasing potency) with increasing treatment duration. (**c)** Model-average BMD analyses and (**d**) confidence intervals in terms of cumulative dose. Here, the BMD confidence intervals stacked nearly on top of one another – demonstrating support for the concept of potency as a function of cumulative dose. (**b/d)** The vertical dashed lines indicate the range of the results by encompassing the lowest-to-highest BMD estimates. In (**d)** the red vertical line indicates the range from just the eight datasets obtained from mice whereas the grey verticals encompass all estimates (*i.e.,* arising from both mouse and rat studies). For each confidence interval, the text indicates the treatment regimen, species, transgene and tissue harvest time-point

Considering the cumulative-dose BMD estimates (**Fig. 5d**) visually, there appears to be a slight trend of increasing BMDs (*i.e.,* lower potency estimations) with increasing exposure durations (*i.e.,* as the regimens move through 1, 4, 9, then finally 28 treatment days the BMDs also increase). This suggests the potential for ‘less-than-additive’ responses with the shift to longer and (typically) lower-dose treatment regimens. However, some of the BMD confidence intervals overlap suggesting this correlation should not be considered statistically significant. Moreover, the two nine-day studies were collected in rat (not mouse) so cross-species sensitivity differences might also contribute to the apparent trend. The benchmark dose analyses underlying **Fig. 5** are presented in **Supplementary Figs. 8-9**.

### Closer scrutiny using error-corrected sequencing

To assess this in more detail, we turned to the recently-published error-corrected next generation sequencing (ecNGS) data obtained from the same liver tissues scored for the TGR endpoint in the GSK study (Ashford et al. 2025). This provided perfectly matched dose-response data for one-day acute versus 28-day, repeat-dose regimens from a technique with unprecedented sensitivity and fidelity that directly quantifies mutations. Using the mutation frequency data from either the ‘Mouse Mutagenesis Panel’ (representative target regions sampled across chromosomes and including coding and non-coding regions) (**Fig. 6a**) or directly from the *lacZ* transgene itself (**Fig. 6b**) yielded BMD estimates on cumulative dose-scales that were *near-identical* (< 1.2-fold different in either panel) (**Fig. 6c**). At the most conservative interpretation (*i.e.,* considering the lowest BMDL to highest BMDU across both treatment regimens) the BMD analyses from the mouse mutagenesis panel suggest that cumulative doses in the range ∼ 2 – 3.6 mg/kg(bodyweight) can be expected to increase mutation frequencies in the mouse liver by 50% (relative to concurrent controls) after daily exposure by gavage for treatment durations between 1 and 28-days in duration. This dose-range was slightly higher for the *lacZ* panel (∼ 4.2 – 6.8 mg/kg(bodyweight). That is, in artificially-inserted bacterial DNA compared to across the mouse genome itself in general. The benchmark dose analyses underlying **Fig. 6** are presented in **Supplementary Fig. 10**.

**Fig. 6-.**
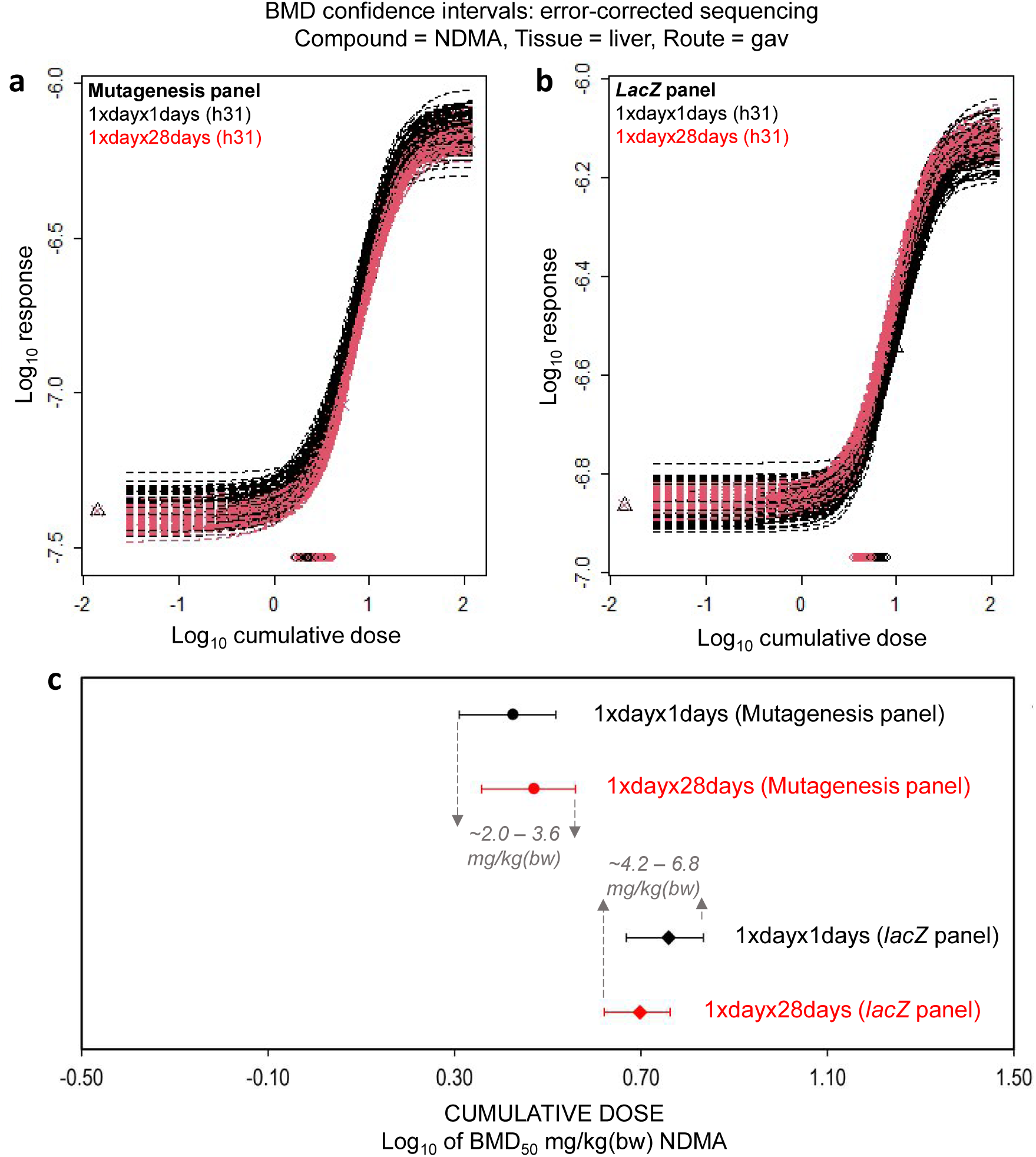
BMD-derived potency comparisons for NDMA using error-corrected sequencing data from the GSK MutaMouse study. (**a**/**b**), Combined-covariate benchmark dose modelling on cumulative dose-scales using mutation frequency dose-response data from either the ‘mouse mutagenesis panel’ or ‘*lacZ* panel’ with regimen as covariate. **c**, BMD confidence interval plots demonstrating near-identical BMDs on cumulative dose scales regardless of single-dose (1xdayx1days) or 28-day, repeat-dose treatment regimens (*i.e.,* 1xdayx28days). The arrows and written dose-ranges indicate the lowest-BMDL-to-highest-BMDU interval across treatment regimens for each target. Both the Mutagenesis panel and *lacZ* panel show excellent support of less-than-lifetime concepts (*i.e.,* mutagenic potency as a function of cumulative dose)

### Supporting evidence from rodent carcinogenicity studies

Having considered the short-term mutagenicity data for NDMA, we wondered if supporting evidence of less-than-lifetime potency relationships could also be found in the available (and longer term) rodent carcinogenicity data. **Table 2** summarises the nine studies found with dose-response data suitable for combined BMD modelling (*i.e.,* at least two-doses plus control). Six datasets were available using matched exposure route (drinking water) with daily treatment durations lasting from 13 to 120 weeks (*i.e.,* up to standard rodent lifetime). Of these six datasets, five had determinable BMDs with tumour manifestation times > 1 year or at lifetime (benchmark dose analyses presented, **Supplementary Fig. 11).** A BMD was not determinable for one of the datasets (Study No. 3) because only two dose-groups were used and a non-monotonic response was observed (*i.e.,* the penultimate dose-group had a higher response than the highest dose-group causing uncertainty in model fitting resulting in an unbounded (infinite) confidence interval) (shown, **Supplementary Fig. 11G)**. Also, a BMDL was not determinable for the 13-week treatment duration study (Study No. 8). This was because only two treatment groups were used with administration by drinking water resulting in similar administered doses of approximately ∼1.24 and ∼1.35 mg/kg(bodyweight)/day. Because both caused quite pronounced effects (animals were 45% and 60% liver tumour-bearing, respectively) the shape of the dose-response curve in the ‘low-dose region’ close to the benchmark response size was uncertain as there were no empirical measurements to anchor it – resulting in an incalculable BMDL (shown, Supplementary Fig. 11C/F).

**Fig. 7a/b** shows the BMD-derived potency estimates in terms of daily-dose. As seen for TGR and in-keeping with less-than-lifetime concepts, the BMDs decreased with increasing treatment duration (*i.e.,* empirically demonstrating increasing carcinogenic potency with increasing treatment duration). Collectively, the BMD estimates occupied a ∼10-fold range. Switching to cumulative dose brought the BMD confidence intervals into much closer agreement with one another (**Fig. 7c/d**). The five BMD estimates now occupied just a ∼2-fold range demonstrating empirical support for the concept of potency as a function of cumulative dose. The BMD analyses suggest that cumulative doses in the range ∼15 – 32 mg/kg(bodyweight) can be expected to increase liver tumour incidence by 10% from control-group levels in rodents after daily exposure via drinking water for treatment durations between 30 weeks and two years in duration. Unfortunately, the results from the 13-week study in mice cannot be included in this summary statement due to the incalculable BMDL.

**Fig. 7-.**
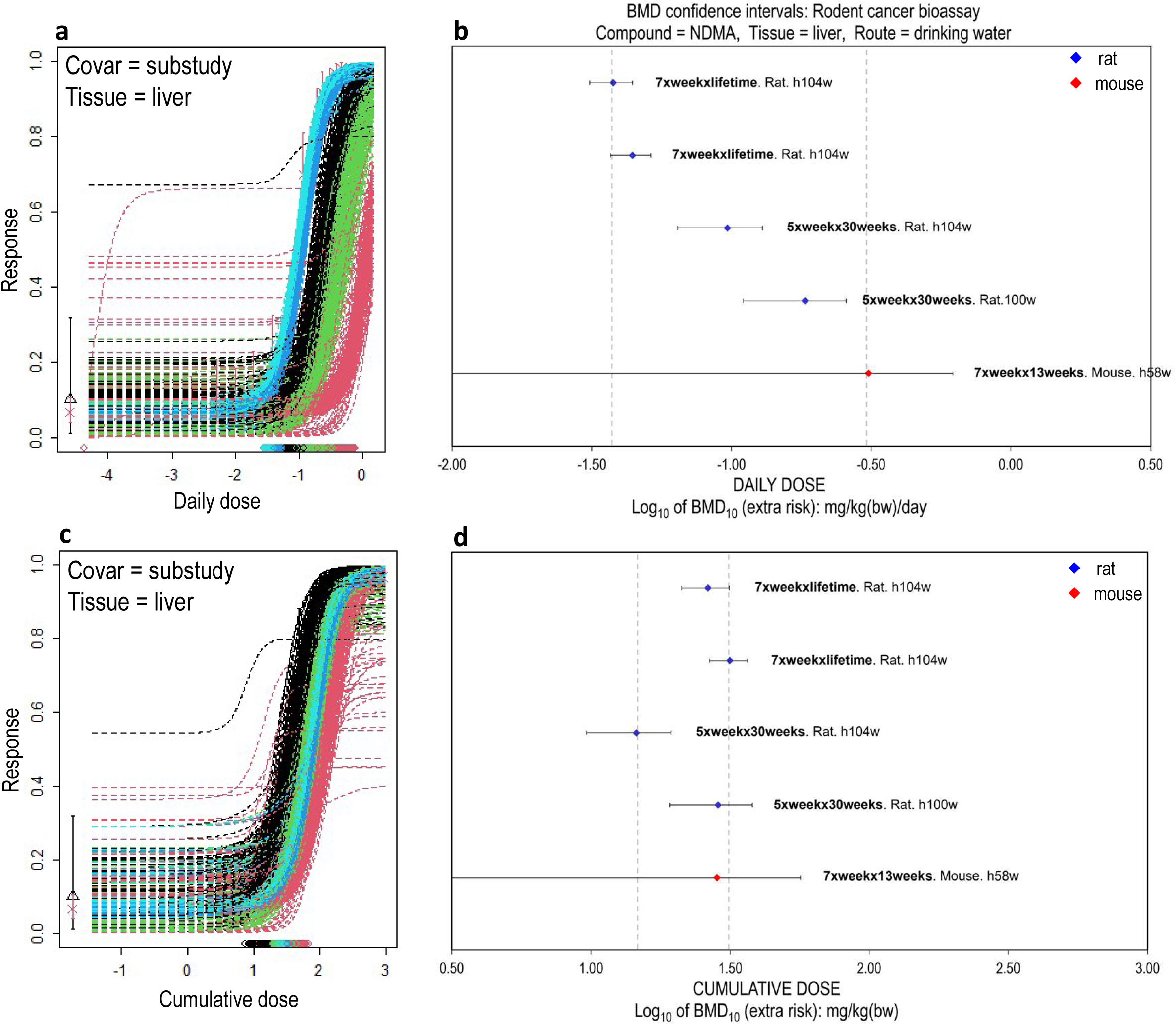
Examining less-than-lifetime potency relationships using rodent carcinogenicity data. (**a**) Model-average BMD analyses and (**b**) resultant confidence intervals on daily-dose scales ranked by increasing treatment duration. (**b**) In-line with less-than-lifetime concepts, BMDs generally decreased (*i.e.,* demonstrating increasing potency) with increasing numbers of treatment weeks. (**c**) BMD analyses and (**d**) confidence intervals calculated using cumulative dose. (**d**) Again, the BMD confidence intervals stack nearly on top of one another – demonstrating support for the concept of potency as a function of cumulative dose. (**b**/**d**) The vertical dashed lines indicate the range of the results by encompassing the lowest-to-highest BMD estimates. BMDL estimates cut-off by the left plot-axis indicate datasets for which a BMDL could not be determined due to uncertainty in the underlying dose-response data. For each confidence interval, the text indicates the treatment regimen, species and tissue harvest time (*i.e.,* the tumour manifestation time).

## DISCUSSION

Understanding less-than-lifetime potency relationships is essential to both the scientific validity and regulatory acceptance of the less-than-lifetime Threshold of Toxicological Concern (TTC) approach as described in ICH M7 (EMA 2023). In practice, (1) most pharmaceuticals are not administered daily over full lifetime, (2) clinical trials rely on limited exposure durations, and (3) achieving and maintaining absolute compound purity is impossible (Felter et al. 2025). Consequently, robust methods for deriving duration-specific acceptable intake limits are critical to effective impurity management, human health risk assessment and successful drug development (Bercu et al. 2021; Felter et al. 2011; Felter et al. 2025).

Previous investigations of less-than-lifetime potency relationships have primarily used TD50 modelling of rodent carcinogenicity data (Thomas et al. 2025). In the present work, BMD modelling was applied as a more precise and statistically robust method for point-of-departure estimation and comparative potency assessment (Thomas et al. 2025; White et al. 2025). In addition to rodent carcinogenicity data, we evaluated durational-potency relationships using dose-response data from shorter-term *in vivo* mutagenicity studies (*i.e.,* transgenic rodent and error-corrected sequencing outcomes) for the data-rich *N-*nitrosamine NDMA.

A recognised challenge in using historical TGR data is the variability in study design (*e.g.,* differences in exposure routes, sampling times, species, strains and transgene reporters *etc.*). This heterogeneity can complicate quantitatively comparable analyses across studies (White et al. 2025; Wills et al. 2017). However, prior work has shown that TGR variant (*i.e.,* species or reporter differences) exert minimal influence on BMD estimates, supporting the broader feasibility of cross-study potency comparisons (Wills et al. 2017). Moreover, covariate-matched historical data were specifically available for NDMA in liver tissue such that we could demonstrate that, provided there was an initial window-of-time for mutation fixation, a broad range of sampling times had no significant effect on the measured BMD.

This again reduced the dependency on exactly-matched study designs and highlighted a difference between the TGR endpoint and the rodent cancer bioassay: whereas TGR responses were largely independent of study duration, cancer bioassay outcomes can vary dramatically due to differences in tumour latency / manifestation time (Bercu et al. 2021; Felter et al. 2025). Collectively, these observations enabled us to harness the globally available TGR and ecNGS datasets for NDMA to test the central tenet of the less-than-lifetime TTC approach – that potency is a function of cumulative dose (EMA 2023).

BMD analyses and point-of-departure estimates are typically conducted and expressed in daily-dose units (*e.g.,* mg/kg(bodyweight)/day). In earlier TGR studies prior to the OECD guideline (OECD 2022) and the more general adoption of 28-day, repeat-dose gavage study designs, key experimental variables included exposure duration and administration route (White et al. 2025; Wills et al. 2017). For risk assessment, such legacy datasets remain valuable, particularly for data-poor compounds, yet daily-dose modelling effectively neglects differences in exposure regimen. Here, for relatively short exposure durations (*i.e.,* ≤ 28 days) BMD estimates for NDMA in liver tissue spanned an ∼80-fold range, whereas cumulative dose scaling compressed this to ∼20-fold and indicated route-specific sub-clusters. When all seven tissues were considered, cumulative-dose analyses more clearly identified the liver as the most sensitive target organ, consistent with the role of CYP2E1 in generating genotoxic NDMA metabolites (Haggerty and Holsapple 1990; Lynch et al. 2024; Yang et al. 1987). Restricting the analysis to the eight most closely matched mouse gavage studies in liver showed that, despite treatment durations from 1 to 28 days, cumulative-dose BMDs varied by only ∼ 2.5-fold. Put another way, the globally-available empirical measurements suggest that cumulative doses of NDMA between ∼ 4 – 10 mg/kg can be expected to cause a 50% increase in mutant frequency responses in mouse livers irrespective of how this total dose is fractionated over the 28 days. The reproducibility of the cumulative-dose estimates derived from studies conducted worldwide over 31 years is striking.

These quantitative analyses provide empirical support for less than lifetime potency relationships for NDMA, a potent small-alkyl *N-*nitrosamine. While mutagenicity data were limited to ≤ 28-day exposures, corresponding carcinogenicity data revealed similar cumulative-dose patterns. Thus, our findings bridge short-term mutagenicity and long-term carcinogenicity through the unifying concept of potency governed by cumulative exposure. Mechanistically, this is expected: although carcinogenesis is a multifactorial process, mutagenesis is a key initiating event shaping the carcinogenic potential of nitrosamines (*i.e.,* driven by alkylation, mutation and therefore increased cancer risk) (Haggerty and Holsapple 1990; Lynch et al. 2024; Yang et al. 1987). Prior studies have reported correlations between *in vivo* mutagenicity (TGR) and rodent carcinogenicity for diverse compounds, including several nitrosamines, further supporting this relationship (Hernández et al. 2011; Jolly et al. 2026; Lambert et al. 2005). Our analyses show that shorter-term mutagenicity assays can reflect the potency relationships observed in longer-term carcinogenicity studies, reinforcing the predictive value of TGR data for cumulative-dose-driven carcinogenicity outcomes.

A frequent concern in applying less than lifetime models to nitrosamines is that short-term, high-dose exposures might overwhelm DNA repair mechanisms, leading to underestimation of risk (Felter et al. 2025; USEPA 1986). Although TGR data beyond 28 days’ exposure are lacking, both ecNGS and TGR datasets demonstrate that single, acute exposures approaching the maximum tolerated dose still yield responses comparable to those cumulated from ‘low dose’ 28-day repeat regimens. Nevertheless, *C* x *t* relationships represent an approximation. Most obviously, high acute doses may be lethal even if the same cumulative dose is tolerable when fractionated, leading to a breakdown of proportionality. Indeed, some TGR studies with NDMA indicate such effects through loss of animals in the highest dose-groups tested (Butterworth et al. 1998; Lynch et al. 2024). Conversely, more efficient elimination in the host, as may occur in the low-dose region due to the efficiency of clearance and compensatory mechanisms may also cause breakdown of *C* inside the limit of *t* such that the assumption of a simple relationship results in potency overestimation. Importantly, in context of ‘low-dose’ impurity management, the former is likely irrelevant whereas the latter suggests the assumption of *C* x *t* may be conservative.

Whereas the TGR analyses suggested such potential for less than additive responses to increasingly longer and lower-dose regimens, analyses of matched samples by ecNGS showed strict conformity to *C* x *t* for both *lacZ* and mouse genome targets for 1 or 28-day treatment durations. This finding does not preclude a role for compensatory mechanisms: it may just be that their subtractive power is similar for both chronic and acute regimens in this instance. It is also noteworthy that the cumulative doses associated with a 50% increase in mutation frequency over control for the *lacZ* panel were ∼ 4 – 7 mg/kg, in strong alignment with TGR (∼ 4 – 10 mg/kg) (Yauk et al. 2025). On cumulative-dose scales the mutagenesis panel was also more sensitive than the *lacZ* panel as indicated by the establishment of non-overlapping BMD confidence intervals. This may reflect differences in target length for the same read-depth and/or adduct suppression due to the heavily methylated and G/C-rich content of the bacterial DNA (Ashford et al. 2025).

Given the shared mechanism of mutagenicity among small alkyl-amine nitrosamines, the less than lifetime potency relationships demonstrated here for NDMA are likely relevant to structurally similar compounds. Indeed, durational potency relationships have been documented for NDEA and other nitrosamines (Bercu et al. 2021; Druckrey 1967; Peto et al. 1991a), as well as for numerous genotoxic carcinogens (Felter et al. 2011). Full development of this work using a gamut of compounds may demonstrate that less than lifetime relationships can be quantitatively demonstrated using shorter-term *in vivo* mutagenicity data – justifying acceptable use of the less than lifetime TTC approach. This work therefore lays the foundations for demonstrating less than lifetime potency relationships using *in vivo* mutagenicity data in place of rodent carcinogenicity outcomes.

Finally, the BMD-based, multi-endpoint approach presented here establishes a route for future case studies with other nitrosamines and chemical classes. Such investigations could enhance understanding of less-than-lifetime potency relationships and the mechanistic continuum from mutagenicity to carcinogenicity. Collectively, these efforts may demonstrate that nitrosamines are not an exceptional ‘cohort of concern’ but behave comparably to other mutagenic carcinogens (Felter et al. 2011; Felter et al. 2025). This would strengthen confidence that the staged TTC approach described in ICH M7 can be appropriately applied to *N-*nitrosamine impurities in both lifetime and less-than-lifetime exposure scenarios.

## SUPPLEMENTARY INFORMATION

Eleven **Supplementary Figures** presenting all of the analyses underpinning the main-text Figures are submitted alongside this Manuscript as an accompanying PDF file.

## ACKNOWLEDGEMENTS

JW would like to thank George E. Johnson for the collaborative brainstorming supporting the analyses presented in this manuscript.

## AUTHOR CONTRIBUTIONS

JWW and AML conceived the study and designed the analyses. JWW, DSGH, JSH and AW collected the data. JWW and RB performed the analyses. JWW wrote the paper in close collaboration with all authors.

## DATA AVAILABILITY

The dose-response data underlying all analyses are available for download from the BioStudies database under accession number S-BSST2655.

## CODE AVAILABILITY

Example R-scripts / the BMD outputs underlying the BMD analyses presented in this work are available for download from the BioStudies database under accession number S-BSST2655.

## CONFLICT OF INTEREST STATEMENT

JWW, AW, DSGH, RB, RH and AML are employees of GlaxoSmithKline.

## ETHICS STATEMENT

All dose-response data unpinning the analyses presented in this manuscript were collected from previously-published literature.

## Notes

https://www.ebi.ac.uk/biostudies/studies/S-BSST2655

### Competing Interest Statement

The authors are employees of pharmaceutical company GlaxoSmithKline.

