## Supplementary Fig for "Mutagenic and carcinogenic potency determinations for NDMA support the cumulative dose assumption underpinning the less-than-lifetime Threshold of Toxicological Concern"

# 20260127
### Supplementary information –

Individual analyses -

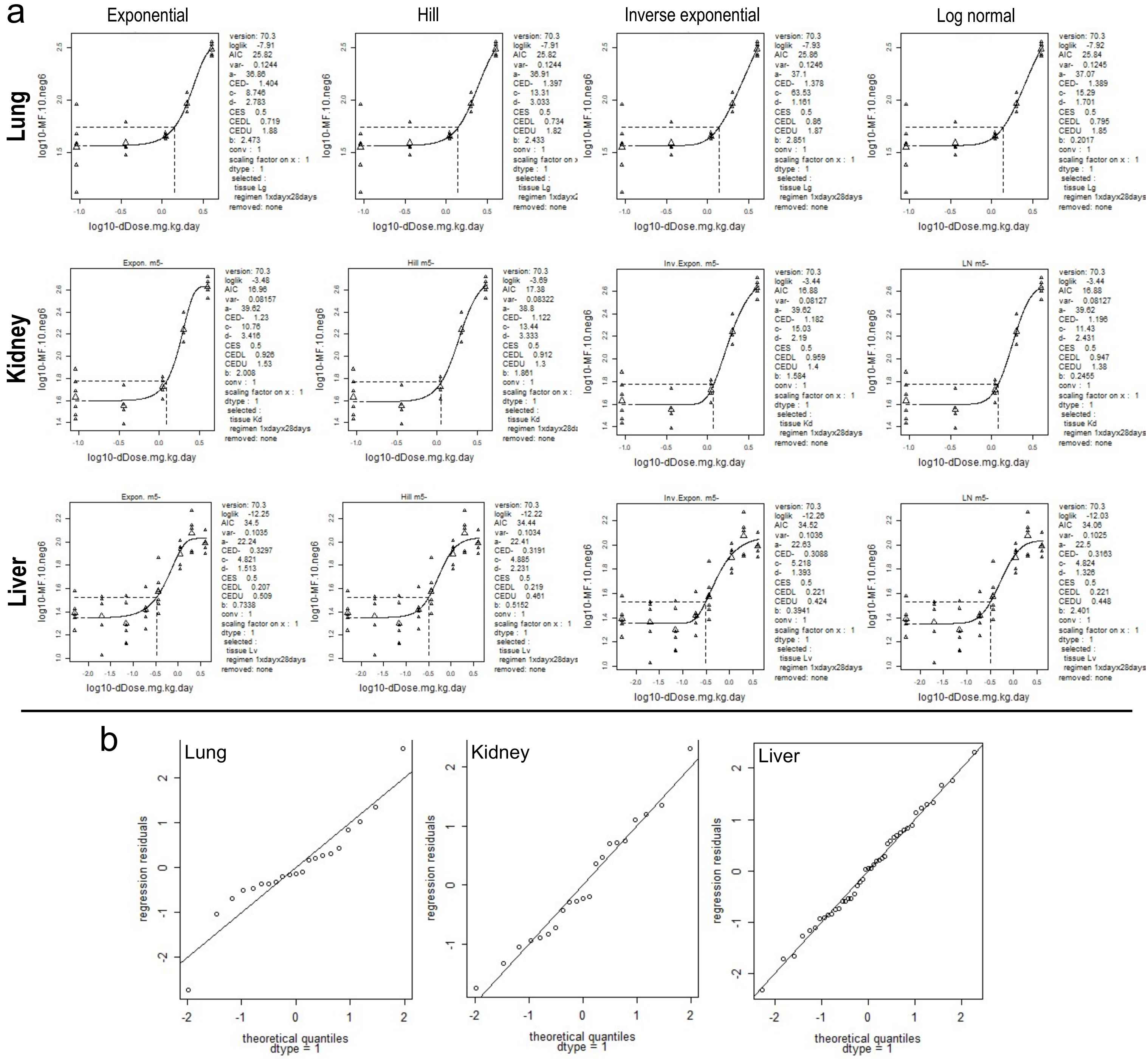


**Suppl. Fig. 1 – Individual BMD analyses behind Figure 2A/B on daily-dose scales.** (**a**) Exponential, Hill, inverse exponential and log-normal model fits to the TGR data collected from lung, kidney or liver tissues. The curves represent the fitted four-parameter models. Horizontal and vertical dashed lines represent the BMR of 50% and BMD50 (respectively). Control-group (i.e., dose zero) response information is placeholdered left-most on each plot given the log10 scale. (**b**) Quantile-quantile plots for exponential model residuals confirming approximate log-normality for each dataset

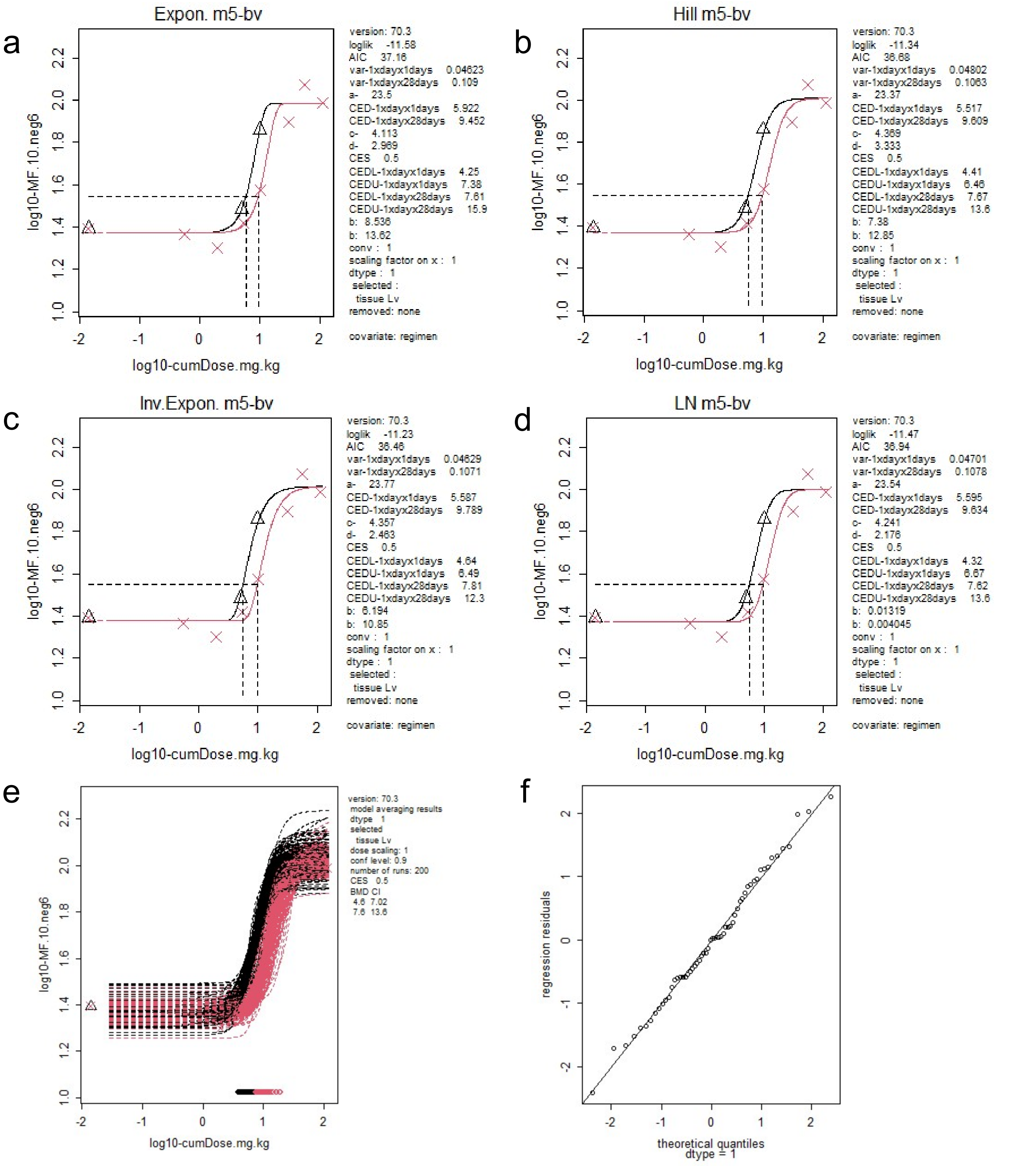


**Suppl. Fig. 2 – BMD analyses behind Figure 2C/D on cumulative-dose scales.** (**a-d**) Exponential, Hill, inverse exponential and log-normal model fits to the TGR data collected from liver tissues with treatment regimen as covariate (red = 1xdayx28days, black = 1xdayx1days). The curves represent the fitted four-parameter models with horizontal and vertical dashed lines indicating the BMR of 50% and BMD50 (respectively). Control-group (i.e., dose zero) response information is placeholdered left-most on each plot given the log10 scale. (**e**) Bootstrap sampling to determine 'model average' BMD confidence intervals. (**f**) Quantile-quantile plots for exponential model residuals confirming approximate log-normality

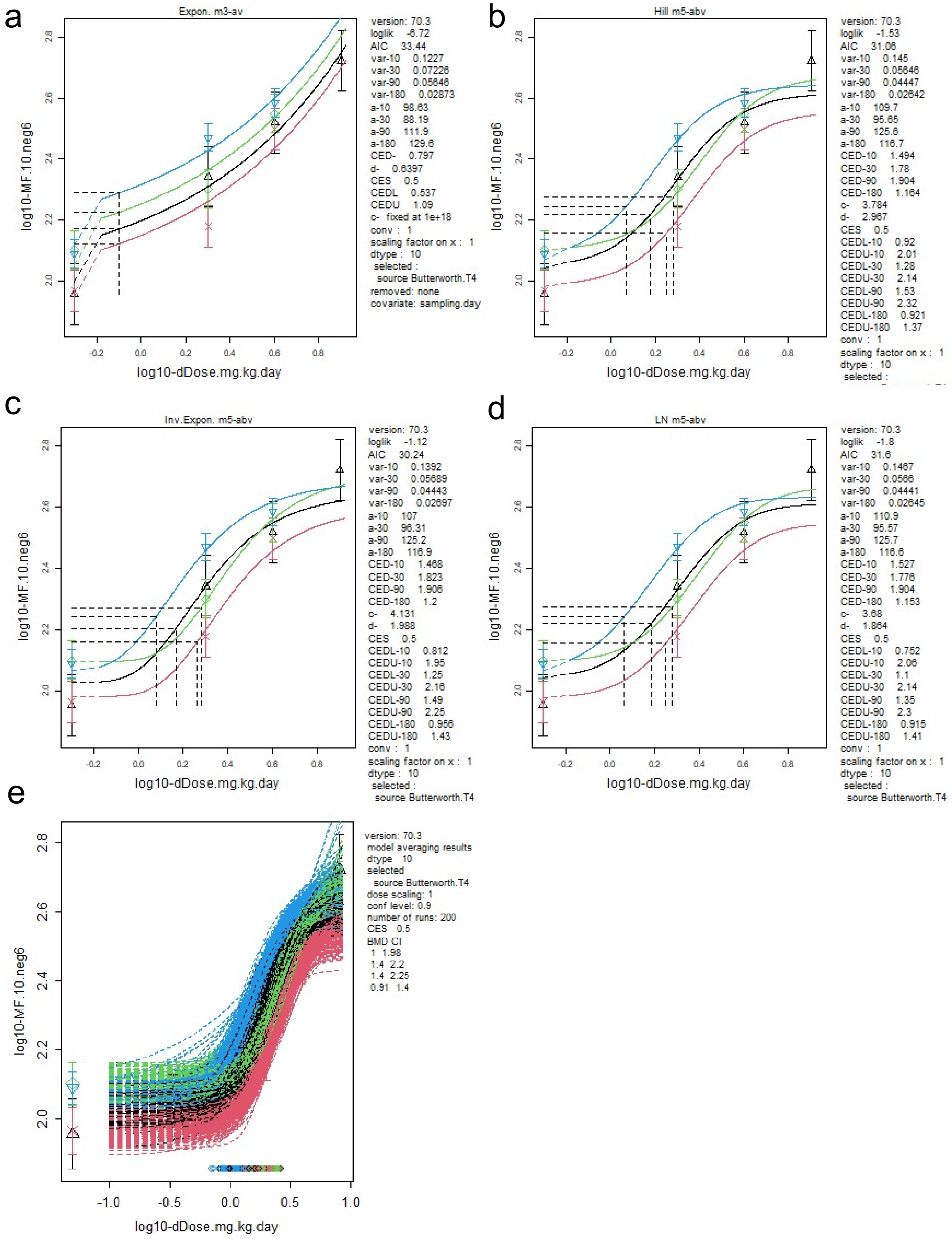


**Suppl. Fig. 3 – BMD modelling behind the Figure 3A/B sampling time analyses on daily-dose scales.** (**a-d**) Exponential, Hill, inverse exponential and log-normal model fits to the summary (i.e., mean response and standard deviation) TGR data collected from liver tissues with sampling time as covariate (blue = 180 days, green = 90 days, black = 10 days, red = 30 days). The curves represent the fitted models with horizontal and vertical dashed lines indicating the BMR of 50% and BMD50 (respectively). Control-group (i.e., dose zero) response information is placeholdered left-most on each plot given the log10 scale. (**e**) Bootstrap sampling to determine 'model average' BMD confidence intervals. Note the (**a**) exponential model family rejects sampling time as an influential covariate and returns a single BMD estimate for all datasets combined across sampling times

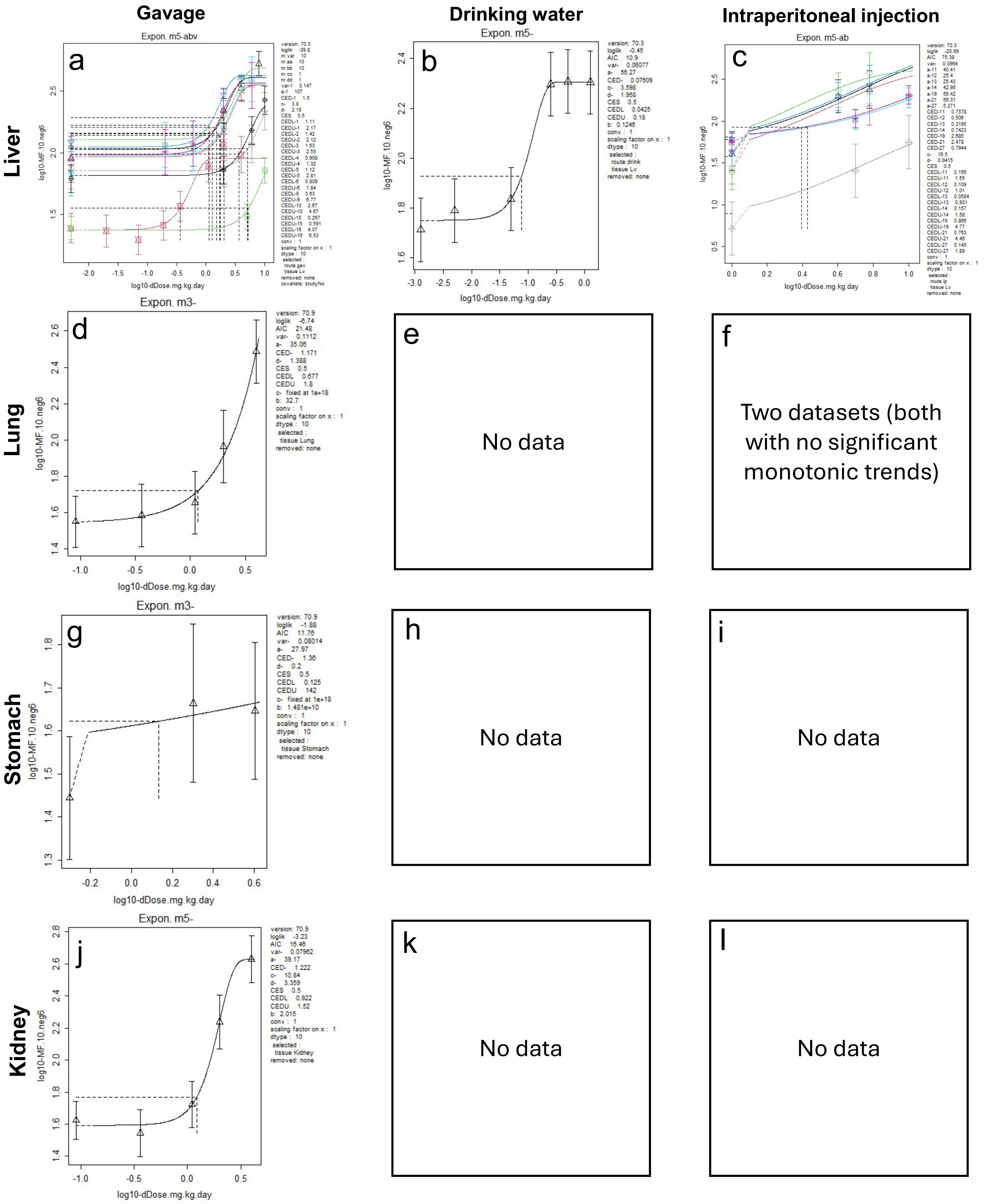


**Suppl. Fig. 4 – BMD analyses behind Figure 4A-D on daily-dose scales.** (**a-l**) Exponential model fits to the summary (*i.e.*, mean response and standard deviation) TGR data collected from the (left) indicated tissues after exposure via the (top) administration routes indicated with 'sub-study' as covariate. The curves represent the fitted model with horizontal and vertical dashed lines indicating the BMR of 50% and BMD50 (respectively). Control-group (*i.e.*, dose zero) response information is placeholdered left-most on each plot given the log10 scale. (**e/f/h/i/k/l**) Absence of plots indicates that either no data were available for the tissue and exposure route combination or (**f**) no monotonic trends were present in the available data

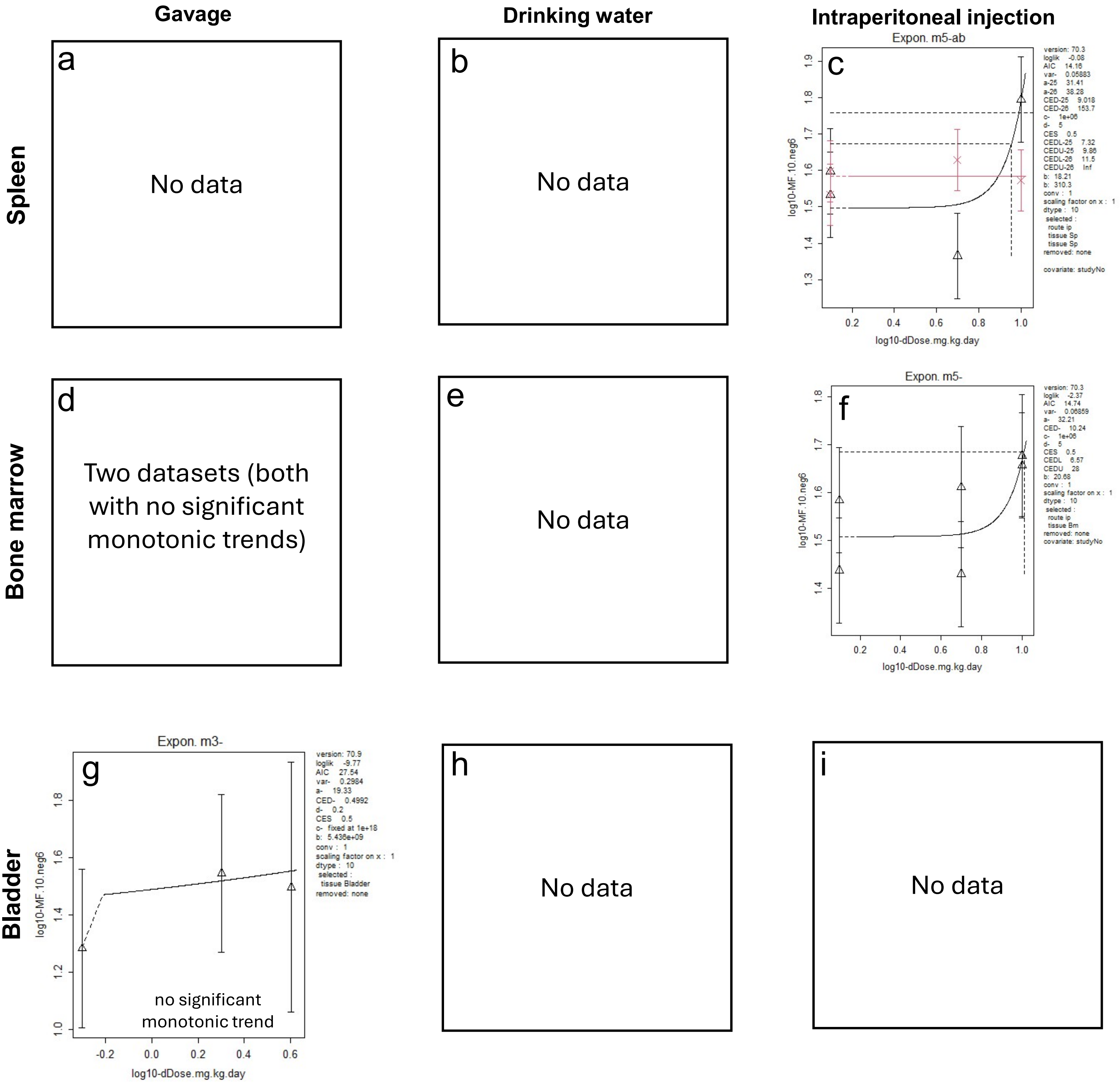


**Suppl. Fig. 5 – BMD analyses behind Figure 4E-G on daily-dose scales.** (**a-i**) Exponential model fits to the summary (*i.e.,* mean response and standard deviation) TGR data collected from the (left) indicated tissues after exposure via the (top) administration routes with 'sub-study' as covariate. The curves represent the fitted model with horizontal and vertical dashed lines indicating the BMR of 50% and BMD50 (respectively). Control-group (*i.e.,* dose zero) response information is placeholdered left-most on each plot given the log10 scale. (**a/b/d/e/h/i**) Absence of plots indicates that either no data were available for the tissue and exposure route combination or (**d**) no monotonic trends were present in the available data

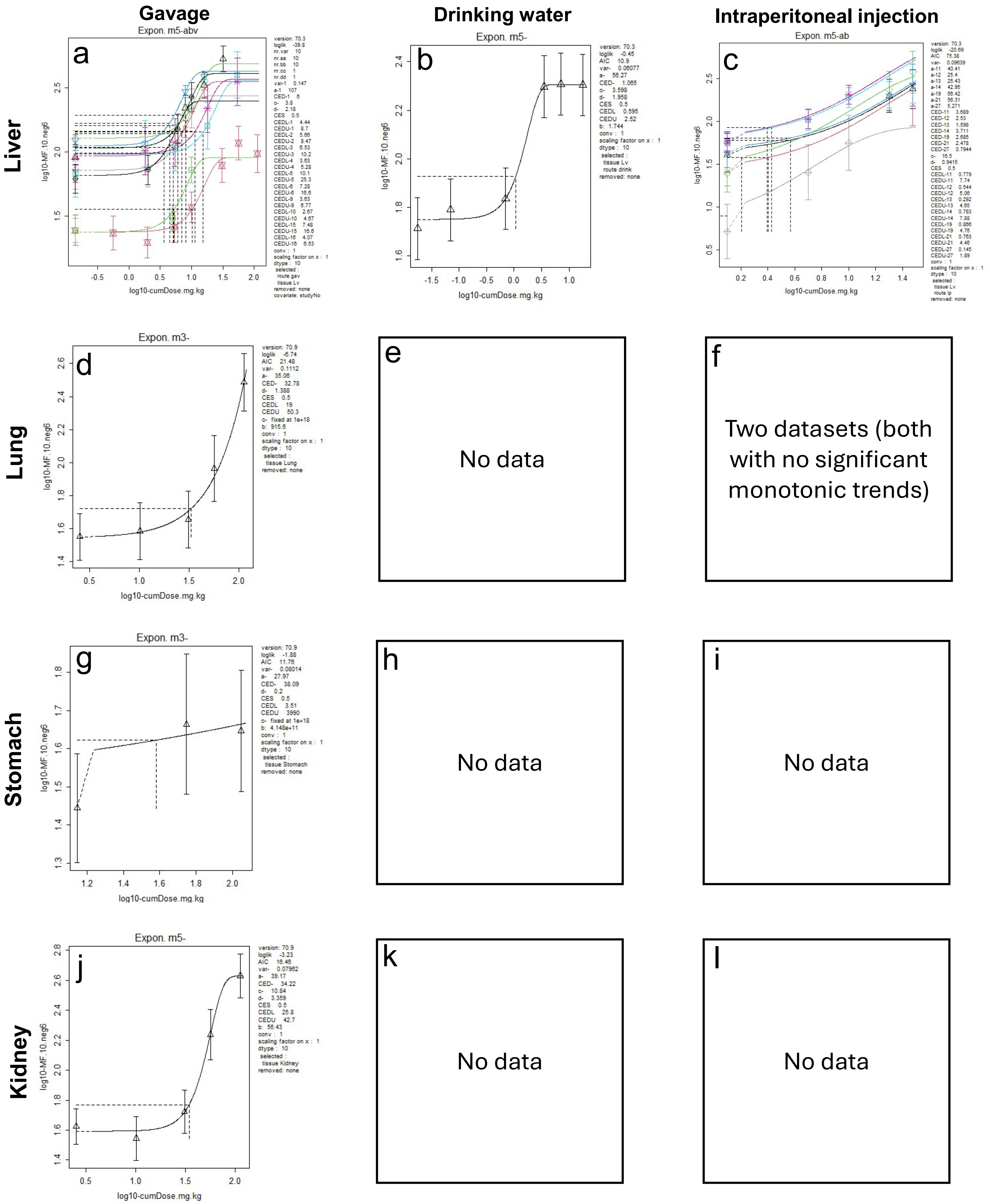


**Suppl. Fig. 6 – BMD analyses behind Figure 4H-K on cumulative-dose scales.** (**a-l**) Exponential model fits to the summary (*i.e.*, mean response and standard deviation) TGR data collected from the (left) indicated tissues after exposure via the (top) administration routes indicated with 'sub-study' as covariate. The curves represent the fitted model with horizontal and vertical dashed lines indicating the BMR of 50% and BMD50 (respectively). Control-group (*i.e.*, dose zero) response information is placeholdered left-most on each plot given the log10 scale. (**e/f/h/i/k/l**) Absence of plots indicates that either no data were available for the tissue and exposure route combination or (**f**) no monotonic trends were present in the available data

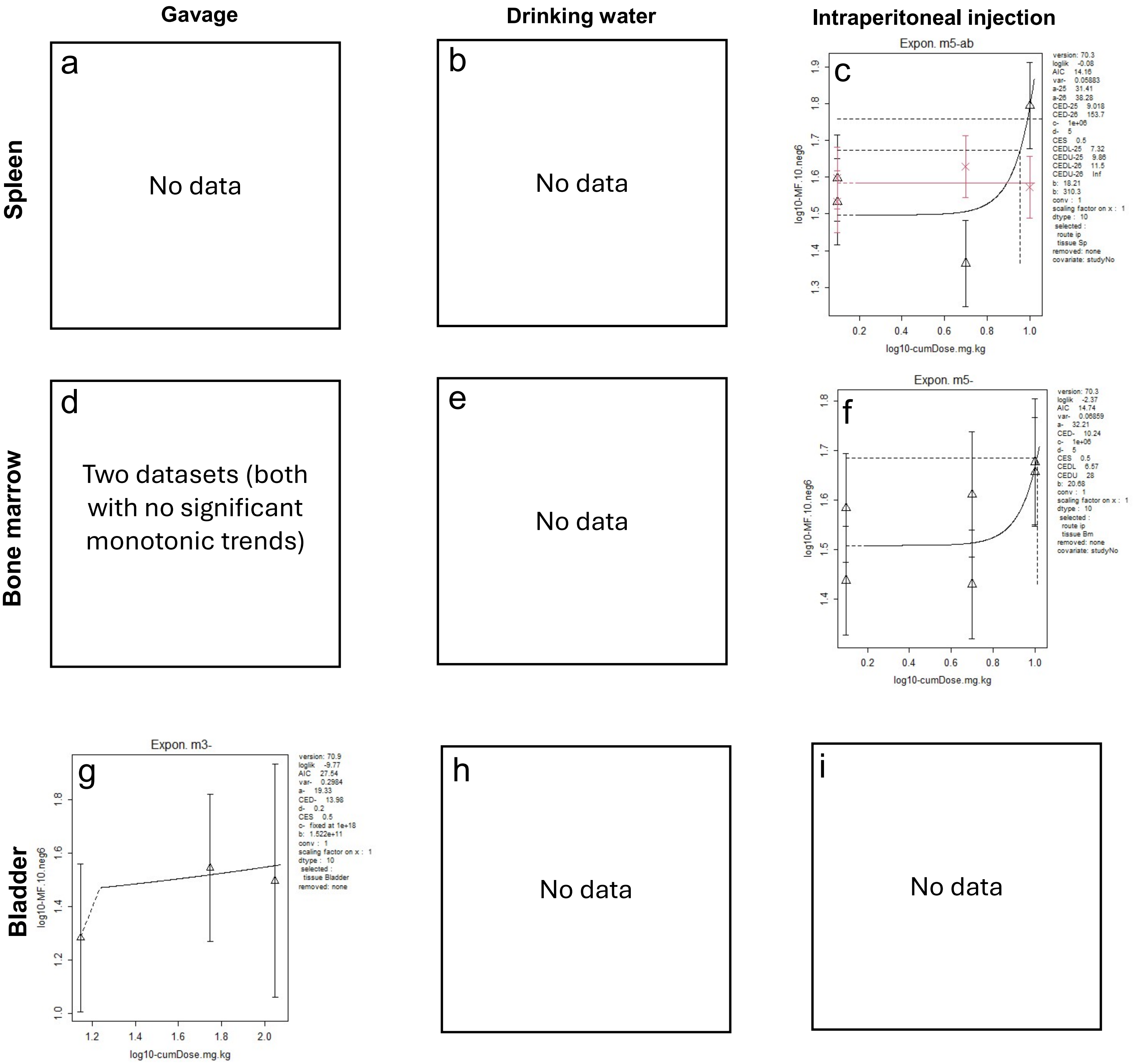


**Suppl. Fig. 7 – BMD analyses behind Figure 4L-N on daily-dose scales.** (**a-i**) Exponential model fits to the summary (*i.e.,* mean response and standard deviation) TGR data collected from the (left) indicated tissues after exposure via the (top) administration routes with 'sub-study' as covariate. The curves represent the fitted model with horizontal and vertical dashed lines indicating the BMR of 50% and BMD50 (respectively). Control-group (*i.e.,* dose zero) response information is placeholdered left-most on each plot given the log10 scale. (**a/b/d/e/h/i**) Absence of plots indicates that either no data were available for the tissue and exposure route combination or (**d**) no monotonic trends were present in the available data

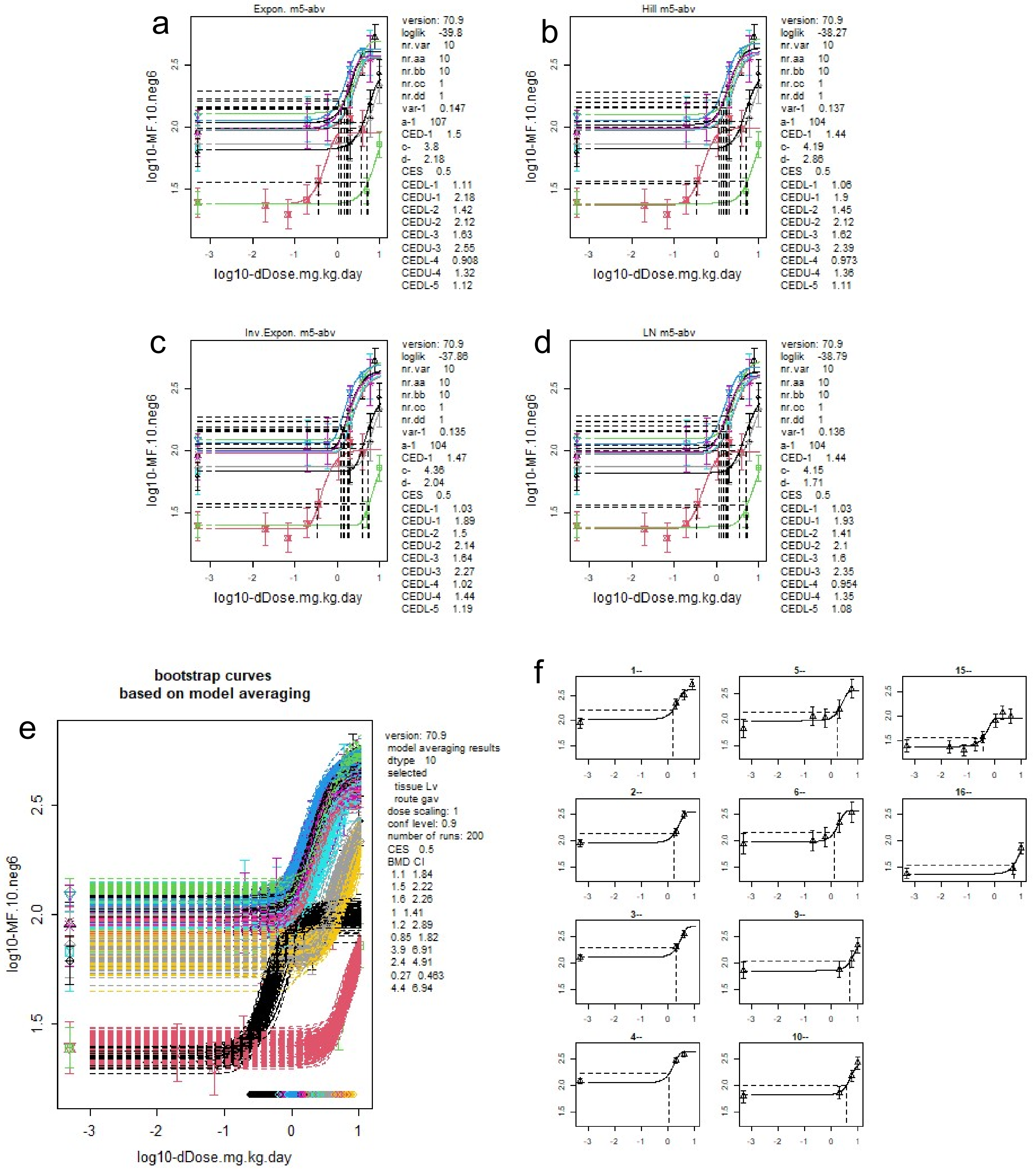


**Suppl. Fig. 8 – BMD analyses behind Figure 5A on daily-dose scales.** (**a-d**) Exponential, Hill, inverse exponential and log-normal model fits to the TGR data collected after gavage administration from liver tissues with 'substudy' as covariate. The curves represent the fitted four-parameter models with horizontal and vertical dashed lines indicating the BMR of 50% and BMD50 (respectively). Control-group (i.e., dose zero) response information is placeholdered left-most on each plot given the log10 scale. (**e**) Bootstrap sampling to determine 'model average' BMD confidence intervals. (**f**) Exponential model fits to each dataset with horizontal and vertical dashed lines indicating the BMR of 50% and BMD50 (respectively)

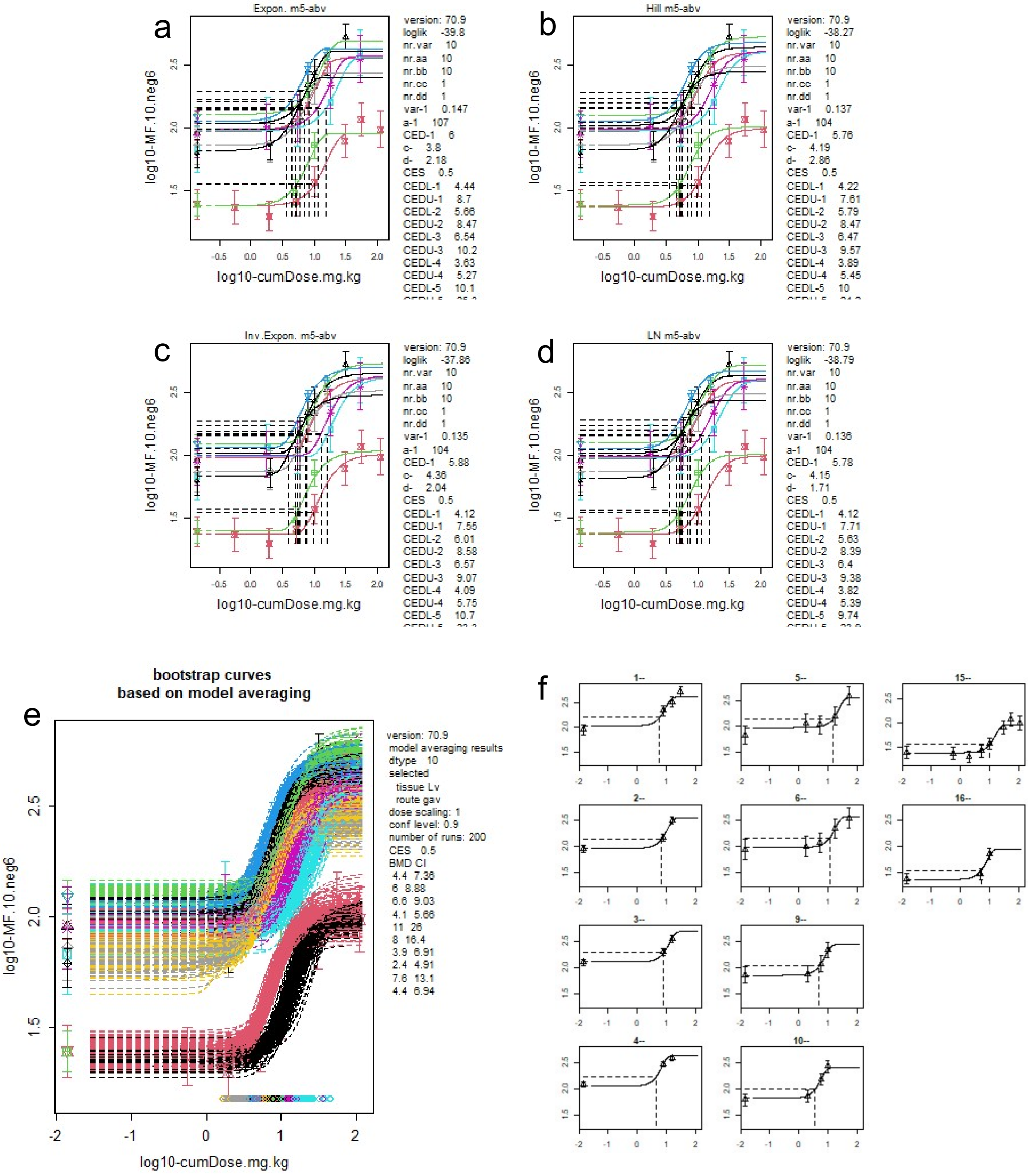


**Suppl. Fig. 9 – BMD analyses behind Figure 5B on cumulative-dose scales.** (**a-d**) Exponential, Hill, inverse exponential and log-normal model fits to the TGR data collected after gavage administration from liver tissues with 'substudy' as covariate. The curves represent the fitted four-parameter models with horizontal and vertical dashed lines indicating the BMR of 50% and BMD50 (respectively). Control-group (i.e., dose zero) response information is placeholdered left-most on each plot given the log10 scale. (**e**) Bootstrap sampling to determine 'model average' BMD confidence intervals. (**f**) Exponential model fits to each dataset with horizontal and vertical dashed lines indicating the BMR of 50% and BMD50 (respectively)

***lacZ* panel -**

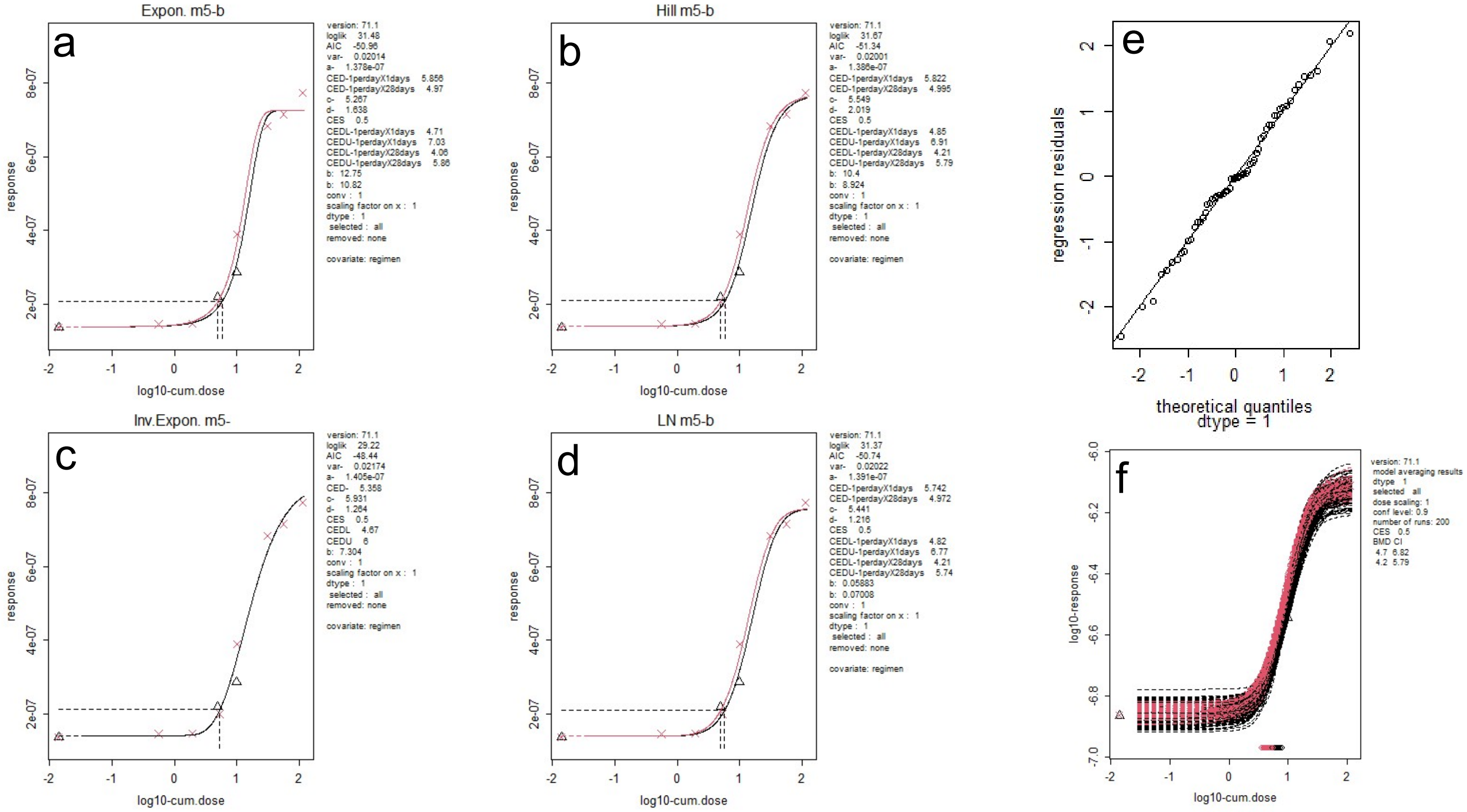


**Mouse mutagenesis panel -**

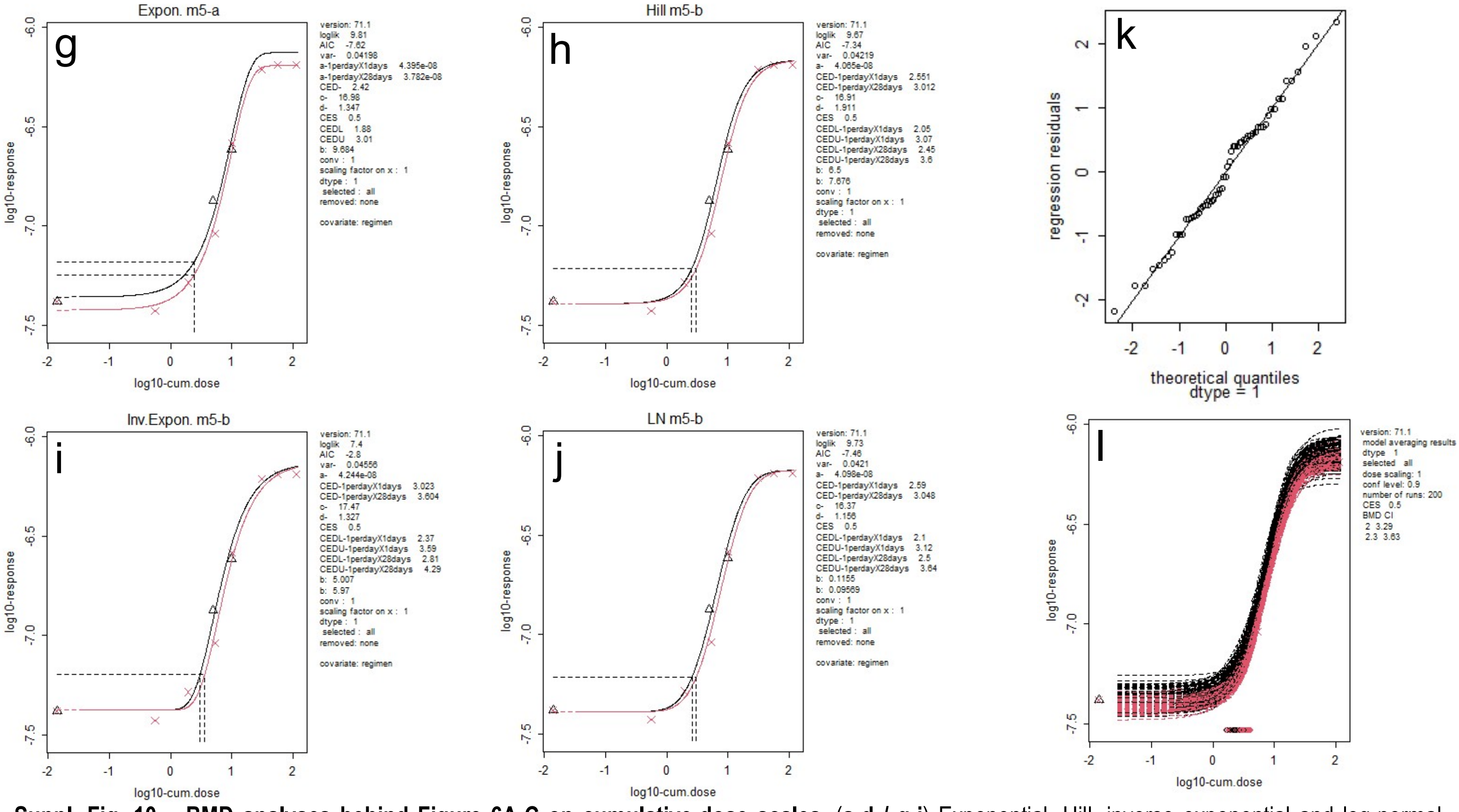


**Suppl. Fig. 10 – BMD analyses behind Figure 6A-C on cumulative-dose scales.** (**a-d / g-j**) Exponential, Hill, inverse exponential and log-normal model fits to the ecNGS mutation frequency data collected from liver tissues with treatment regimen as covariate (red = 1xdayx28days, black = 1xdayx1days). Top shows outputs from the '*lacZ* panel' whilst bottom shows outputs from the 'mouse mutagenesis panel'. The curves represent the fitted four-parameter models with horizontal and vertical dashed lines indicating the BMR of 50% and BMD50 (respectively). Control-group (i.e., dose zero) response information is placeholdered left-most on each plot given the log10 scale. (**e/k**) Quantile-quantile plots for exponential model residuals confirming approximate log-normality. (**f/i**) Bootstrap sampling to determine 'model average' BMD confidence intervals. (**f**) Exponential model fits to each dataset with horizontal and vertical dashed lines indicating the BMR of 50% and BMD50 (respectively)

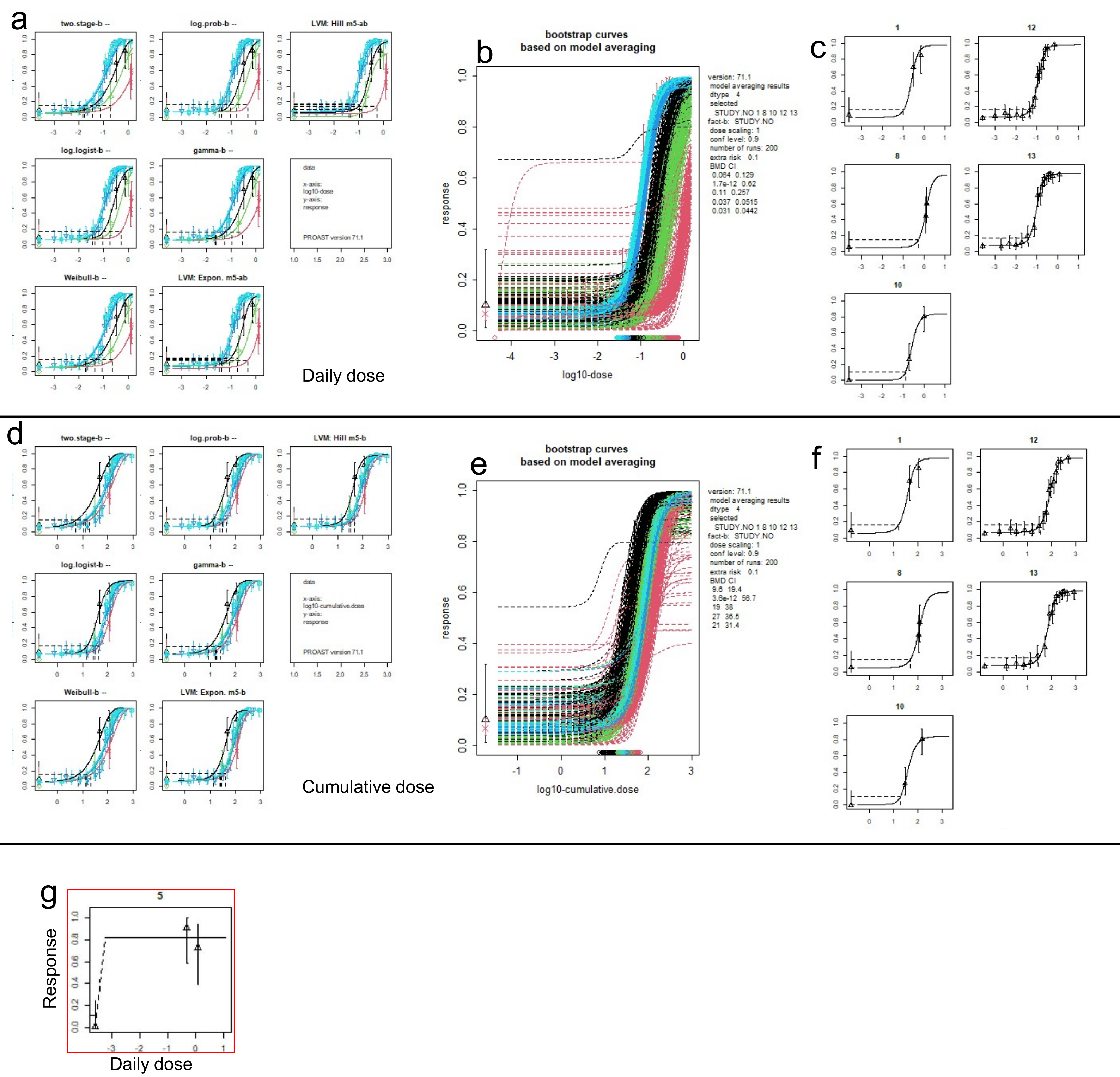


**Suppl. Fig. 11 – BMD analyses behind Figure 7A-D on (a-c) daily-dose or (d-f) cumulative-dose scales.** (**a/d**) Two-stage, log-probit, Hill, log-logistic, gamma, Weibull and exponential model fits to the rodent cancer bioassay data collected after drinking water administration from liver tissues with 'substudy' as covariate. The curves represent the fitted models with horizontal and vertical dashed lines indicating the BMR of 10% (extra risk) and BMD10 (respectively). Control-group (i.e., dose zero) response information is placeholdered left-most on each plot given the log10 scale. (b/e), Bootstrap sampling to determine 'model average' BMD confidence intervals. (**c/f**), Log-logistic model fits to each dataset with horizontal and vertical dashed lines indicating the BMR of 10% and BMD10 (respectively). **g**, One dataset was removed from model fitting as the non-monotonic nature of the response data caused the PROAST fitting algorithm to crash
